# A non-catalytic role of TET3 promotes open chromatin and enhances global transcription

**DOI:** 10.1101/177626

**Authors:** Christel Krueger, Julian R. Peat, Melanie A. Eckersley-Maslin, Timothy A. Hore, Hisham Mohammed, Simon R. Andrews, Wendy Dean, Wolf Reik

## Abstract

The methylcytosine dioxygenase Tet3 is highly expressed as a specific isoform in oocytes and zygotes but essentially absent from later stages of mouse preimplantation development. Here, we show that Tet3 expression promotes transdifferentiation of embryonic stem cells to trophoblast-like stem cells. By genome-wide analyses we demonstrate that TET3 associates with and co-occupies chromatin with RNA Polymerase II. Tet3 expression induces a global increase of transcription and total RNA levels, and its presence further enhances chromatin accessibility in regions open for transcription. This novel function of TET3 is not specific to the oocyte isoform, independent of its catalytic activity, the CXXC domain, or its interaction with OGT, and is localised in its highly conserved exon 4. We propose a more general role for TET3 promoting open chromatin and enhancing global transcription during changes of cellular identity, separate from its catalytic function.

## Introduction

The Ten-eleven translocation methylcytosine dioxygenase (TET) family of proteins catalyse the oxidation of 5-methylcytosine (5mC) to 5-hydroxymethylcytosine (5hmC) and further oxidation products (Tahiliani et al. 2009; Ito et al. 2010, 2011; He et al. 2011). Dilution or removal of these modified bases is one mechanism by which DNA methylation can be reset in the genome (Branco et al. 2012; Kohli and Zhang 2013; Rasmussen and Helin 2016). In addition to the catalytic domain, TET1 and TET3 encode a CXXC domain, which is proposed to aid in targeting the enzymes to specific genomic regions (Xu et al. 2011, 2012; Long et al. 2013). While less studied, TET proteins also have non-catalytic roles, in particular by recruiting other chromatin and epigenetic modifiers, most notably O-linked N-acetylglucosamine transferase (OGT), to their genomic targets (Vella et al. 2013; Chen et al. 2013; Deplus et al. 2013; Ito et al. 2014).

Of the three family members, Tet3 is the least understood, partially due to its unique expression pattern. In contrast to Tet1 and Tet2, Tet3 is robustly expressed in mouse oocytes and zygotes, after which it is rapidly silenced and largely absent by the 4-cell stage (Gu et al. 2011; Iqbal et al. 2011; Wossidlo et al. 2011), coinciding with overall degradation of maternal RNA (Li et al. 2010, Fig. EV1A). In the current model, TET3 is responsible in part for demethylation of the paternal genome and also contributes to the removal of 5mC in the maternal pronucleus (Gu et al. 2011; Iqbal et al. 2011; Wossidlo et al. 2011; Santos et al. 2013; Guo et al. 2014; Peat et al. 2014; Shen et al. 2014). However, this view has recently been challenged: Amouroux and co-workers argue that initial loss of paternal 5mC is mechanistically uncoupled from 5hmC formation (Amouroux et al. 2016). Given TET3’s striking expression pattern and its proposed role in the resetting of epigenetic marks in the early embryo, it was rather unexpected that lack of TET3’s catalytic function was compatible with pre-implantation development (Gu et al. 2011; Peat et al. 2014; Tsukada et al. 2015). We were therefore curious to investigate potential other functions of TET3 unrelated to DNA demethylation.

Here, through integration of transcriptome analysis, assessment of chromatin accessibility and genome-wide chromatin-immunoprecipitation, together with proteomic analysis and global transcription/RNA assays, we provide detailed evidence that TET3 induces hypertranscription and open chromatin independent of its catalytic function. We discuss these results in the light of a role for TET3 in facilitating changes of cell identity.

## Results

### Tet3 is expressed during cellular transitions and promotes change of cell identity

Oocytes produce very high levels of Tet3 (Gu et al. 2011; Iqbal et al. 2011; Wossidlo et al. 2011). Analysis of RNA-seq data from oocytes (Smallwood et al. 2011) and quantitative reverse-transcription PCR indicated the existence of an oocyte specific promoter and a characteristic splice isoform of Tet3. By analysing deep RNA-sequencing (RNA-seq) data of mouse oocytes (Veselovska et al. 2015), we found that the Tet3 transcript produced in oocytes indeed originates from an upstream promoter which adds a small oocyte specific exon and mostly omits exon 2 which encodes the CXXC domain (Fig. 1A) yet retains the catalytic domain. The promoters upstream of exon 2 and exon 4 which are active in other tissues are not used in oocytes. Supporting our analysis, an oocyte specific Tet3 isoform has also recently been reported by Jin et al. (2016).

**Figure 1.**
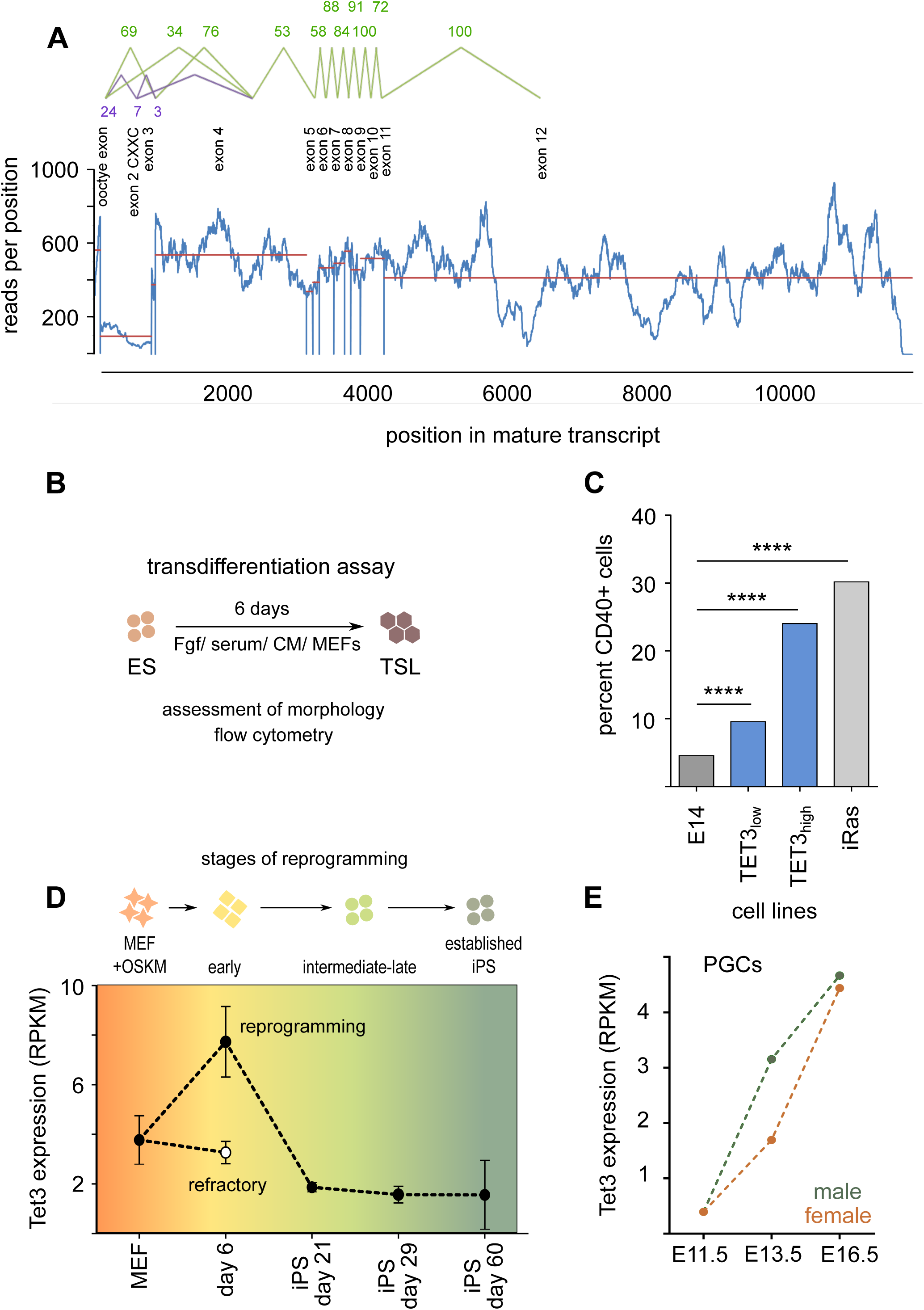
A) TET3 exon usage in germinal vesicle oocytes (data from (Veselovska et al. 2015)). RNA-seq reads were quantified per position in transcript and are shown in blue. Averages over exons are shown in red. Splice junctions were quantified as percentages of the most abundant splice junction and are shown in green for the oocyte specific transcript (Tet3) and in purple for the CXXC containing transcript (Tet3_CXXC_). The majority of transcripts start at the oocyte promoter and most are spliced directly onto exon 3 skipping the CXXC domain containing exon 2. Splicing from the oocyte exon onto exon 2 results in a frame shift. B) Experimental setup of the transdifferentiation assay: embryonic stem (ES) cell lines are cultured in trophoblast stem (TS) cell conditions (see methods) for 6 days. TS-like (TSL) cells are identified morphologically and by immunofluorescent staining for the trophoblast marker CD40. C) Percentage of CD40 positive cells at day 6 of transdifferentiation. iRas cells were used as a positive control for facilitated transdifferentiation (Cambuli et al. 2014). E14 ES cells were used as a negative control. TET3_low_ and TET3_high_ denote clonal cell lines with low and high expression levels of Tet3 respectively. **** p<0.0001, Chi-squared test. D) Tet3 expression during somatic reprogramming monitored by RNA-seq (data from (Milagre et al. 2017). MEF: mouse embryonic fibroblasts; OSKM: ectopic expression of Oct4, Sox2, Klf4 and cMyc; iPS: induced pluripotent stem cells; reprogramming cells are negative for Thy1 and positive for SSEA1; refractory cells are positive for Thy1 and negative for SSEA1 (Polo et al. 2012). E) Tet3 expression during primordial germ cell (PGC) development monitored by RNA-seq (data from (Seisenberger et al. 2012). E: embryonic day. Green datapoints represent levels in male embryos, orange datapoints levels in female embryos.

This led us to hypothesise that the oocyte specific isoform of Tet3 may have a different function to somatically expressed Tet3. Since its expression window coincides with the establishment of totipotency *in vivo*, we explored the possibility that expression of Tet3 is involved in the fundamental reorganisation from gametes to zygote and potentially enhances cellular potency. To test this, we ectopically expressed the oocyte specific isoform of Tet3 (Tet3 for brevity) in ES cells which normally express only very low levels of the gene (Wossidlo et al. 2011) and analysed their potential to transdifferentiate into trophoblast stem-like (TSL) cells.

ES cells are derived from the inner cell mass of the blastocyst at a developmental stage at which the first lineage segregation has already occurred: the inner cell mass is the precursor of all embryonic lineages but cannot contribute to the extra-embryonic lineages as a firm lineage barrier has already been established (Rossant 2008). Analogously, in vitro, ES cells do not readily differentiate into their extra-embryonic counterpart, trophoblast stem (TS) cells (Ng et al. 2008). However, trans-differentiation assays, which employ defined culture conditions, can coax ES cells into resembling TS cells (TS-like cells), albeit at very low frequency (Fig. 1B, C). Transdifferentiation of ES into TS-like cells can be boosted by DNA demethylation (Ng et al. 2008), the ablation of embryonic pluripotency factors such as Oct4, or ectopic overexpression of early trophoblast transcription factors including Cdx2 (Cambuli et al. 2014). We compared the transdifferentiation ability of TET3 expressing ES cells to control ES cells (E14) and well characterised transdifferentiation models (constitutive activation of the Ras ATPase or Oct4 knock-out (Cambuli et al. 2014)). In contrast to control ES cells, many colonies of TET3 expressing cells subjected to transdifferentiation conditions displayed a flat, epithelial-like morphology reminiscent of TS cells (Fig. EV1B). Additionally, the percentage of cells expressing the surface marker CD40 was substantially elevated in Tet3 overexpressing ES cells (24 % compared to 5 %), nearly reaching the level of the Ras-induced transdifferentiation (30 %, Figure 1C). The cell surface marker CD40 is expressed in TS cells, but not in ES cells (Rugg-Gunn et al. 2012). Of note, the ability of cells to acquire features of TS-like cells was dependent on the level of TET3, with Tet3 high-expressing cells moving more towards the trophoblast lineage than Tet3 low-expressing cells (Figure 1C, EV1C).

Interestingly, Tet3 expression is very low in ES cells (Wossidlo et al. 2011) and not substantially upregulated in TS cells (expression data from Adachi et al. 2013). TET3 is therefore unlikely to induce expression of a lineage-specific set of genes. Thus, we interpret the finding that TET3 enhances transdifferentiation from ES to TS-like cells as an ability to expand cellular potency and/or facilitate changes of cell identity.

Intriguingly, apart from its roles in early embryonic development TET3 expression changes have also been reported in a number of seemingly unrelated studies: For example, while Tet3 levels are low in both ES and epiblast stem (EpiES) cells, its transcript is temporarily upregulated during differentiation from ES to EpiES cells (Veillard et al. 2014). Moreover, Tet3 is higher in embryos produced by somatic cell nuclear transfer compared to *in vitro* fertilisation in bovine (Hosseini et al. 2016), relating its upregulation to somatic reprogramming. Also, its expression was found to be beneficial for intracellular sperm injection (ICSI) outcome in humans (Ni et al. 2016). The unifying theme of these reports is the presence and potential role of Tet3 at transition stages.

Extending these observations, we monitored Tet3 expression in an experimental system in which not only the endpoints but also intermediate stages are well characterised, namely reprogramming from mouse embryonic fibroblasts (MEFs) to induced pluripotent stem (iPS) cells. Using data from (Milagre et al. 2017) we found that TET3 is strongly upregulated during early reprogramming (day 6) but almost absent in late stages (day 21, 29) and fully reprogrammed iPS cells (Fig. 1D). Importantly, at day 6, only cells that will go on to be reprogrammed express high levels of Tet3, but those that prove refractory do not, suggesting that the peak in Tet3 expression may be required for successful initiation of reprogramming.

To further explore a potential role in cell identity transitions, we investigated Tet3 levels during primordial germ cell (PGC) development *in vivo* (data from Seisenberger et al. 2012). Tet3 transcript levels increase substantially from E11.5 to E16.5 (Fig. 1E). Of note, this occurs in both male and female PGCs, arguing against an early build-up of transcript for the high levels observed in the mature oocyte.

Since these examples of Tet3 expression coincide with a change, but not necessarily an increase, in cellular potency, and the molecular analysis of different Tet3 isoforms did not support a unique function for oocyte specific TET3 (see below), we propose that TET3 can facilitate changes in cell identity. The transition from highly specialised games to totipotent zygote would present an extreme case of reworking cellular identity, and is accompanied by the highest Tet3 expression levels.

### TET3 expression increases transcription and global RNA levels

We next analysed whether the increased ability to change cell fate was a result of specific transcriptional changes. For this we performed total RNA-seq on TET3 overexpressing ES and control cells (E14 stably transfected with a Tet3 expression construct or empty vector, see Methods and Table S6). Tet3 expression levels were similar to expression levels in oocytes (Figure EV2). To our initial surprise only nine genes were robustly differentially expressed (Fig. 2A, Table S1; for analysis and interpretation of differentially expressed genes using less stringent filtering see Text S1, Fig. S1). No gene ontology groups or biological pathways were prominent, and we could not infer any biological significance of the differentially expressed genes. Interestingly, analysis of the repetitive fraction of the transcriptome showed a dramatic upregulation of MERVL endogenous retroviral elements (Fig. 2B). MERVL expression is characteristic for the transcriptome of 2-cell embryos (Kigami et al. 2003; Evsikov et al. 2004; Peaston et al. 2004) and drives a network of early embryonic genes (Macfarlan et al. 2011). While genes part of this network are generally upregulated in Tet3 overexpressing cells, they do not pass stringent thresholds for differential expression. MERVL elements are also expressed during iPSC reprogramming at a time when Tet3 expression peaks (Eckersley-Maslin et al. 2016).

**Figure 2.**
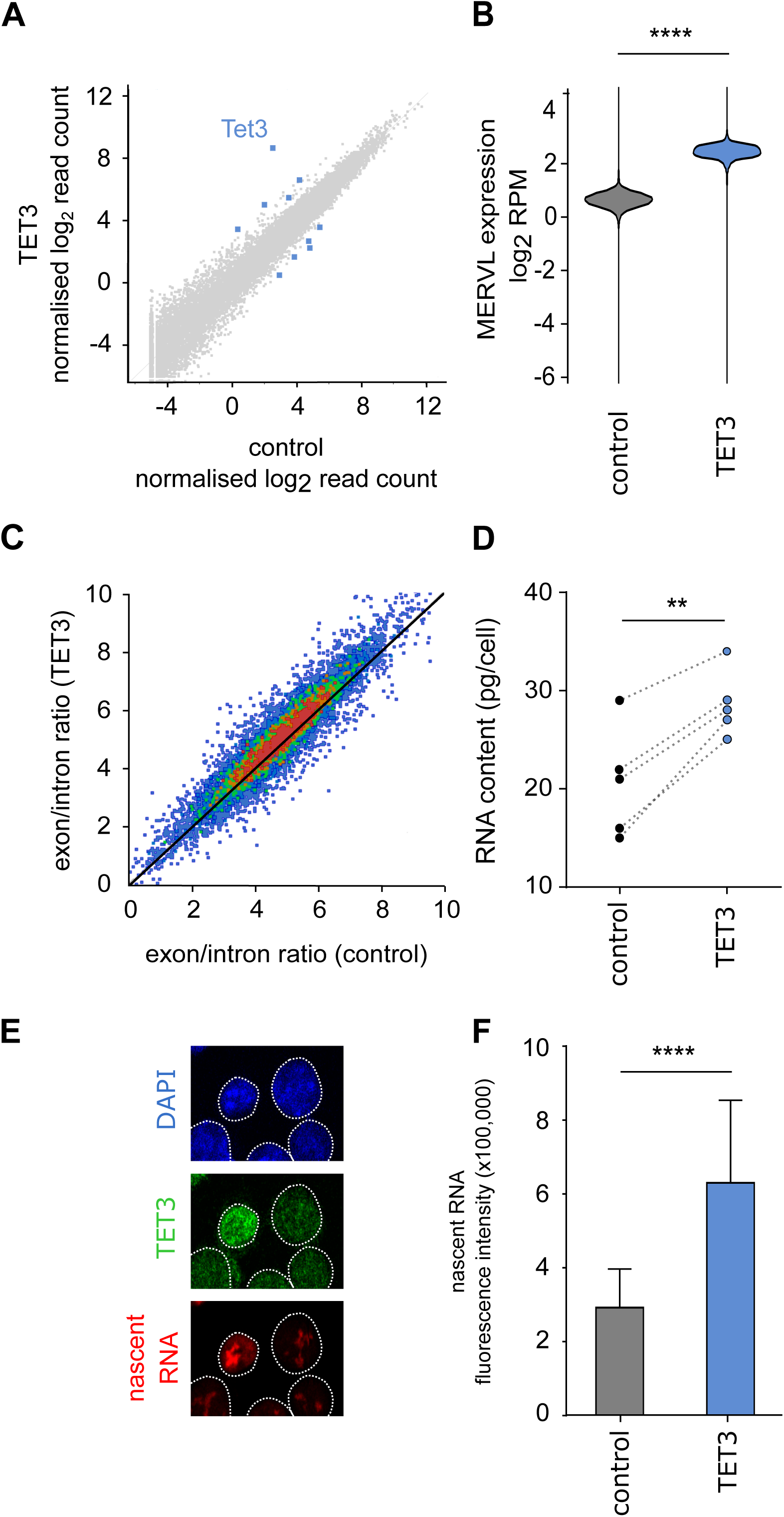
A) Scatter plot showing total RNA-seq data quantified as log_2_ read count per mRNA normalised to the total number of reads. Every dot represents one transcript. Blue dots: Genes differentially expressed between TET3 expressing and control cells (intersection of DESeq2 and intensity difference filters, see methods). Expression level of Tet3 is denoted. B) Bean plot showing expression of full-length MERVL elements in TET3 overexpressing (TET3) and control cells. The LTR ERVL annotation generated by RepeatMasker was filtered for elements longer than 5 kb and RNA-Seq data quantitated as log_2_ RPM. **** p < 0.0001, Mann-Whitney test. C) Density scatter plot showing ratios of exonic over intronic reads obtained for genes with at least minimum expression (log_2_ RPM > -2). Data is from total RNA-seq depleted for ribosomal RNA (rather than PolyA enriched) with on average around 75 % of genic reads originating from exons and 25 % of genic reads originating from introns. D) RNA was isolated from defined cell numbers (116 – 200 × 10^3^ cells) and assessed using fluorometric quantitation. Five independent experiments are shown and data pairs indicated by dashed lines. ** p=0.0025, two-tailed paired t-test. E) Fluorescent labelling of TET3 and nascent RNA. Images show example cells with higher and lower expression of TET3 (middle panel, green) and newly transcribed RNA (bottom panel, red). Nuclei were identified by DAPI staining (top panel, blue) and white dotted lines depict the nuclear outlines. F) Quantification of fluorescent labelling in E. Bars show average and standard deviation of nascent RNA signal per cell in TET3 positive (n=154) and control cells (n=310). **** p<0.0001, Mann-Whitney test.

Strikingly, however, we noticed a globally increased exon to intron ratio in TET3 expressing cells suggesting that there was overall more mature relative to nascent transcript (Fig. 2C). Amongst other possibilities, a higher exon/intron ratio would be observed in RNA-seq data if the cells contained more RNA in total since mature transcripts comprise the bulk of the total read count to which samples are normalised. This could be due to a global change in transcription, RNA processing and/or turnover. Unfortunately, since we had not anticipated a global effect, our experimental setup did not feature controlled cell numbers nor external references which made it impossible to discriminate between these possibilities (Percharde et al. 2017b). However, the finding prompted us to investigate these further.

We next carefully measured the RNA content in a defined number of cells using sensitive fluorimetric quantitation. Indeed, we found that TET3 expressing cells contained 1.5-fold more total RNA than control cells (29 pg/cell vs. 20 pg/cell, Fig. 2D). Additionally, to directly measure transcriptional output of TET3 overexpressing versus control cells and distinguish between an increase in transcription rate and RNA processing and/or turnover, we monitored the production of nascent RNA using a fluorescence based assay (Jao and Salic 2008). Importantly, TET3 overexpressing cells showed 2-fold higher signals for nascent RNA than control cells (Fig. 2E, F; specificity of antibody Fig. S2A). Taken together, data from RNA-seq, nascent transcription assays and quantification of total RNA demonstrate that TET3 substantially increases transcriptional output and raises the RNA content per cell.

### TET3 binds to chromatin regions occupied by RNA Polymerase II

We next asked which genomic regions were bound by TET3 using chromatin immunoprecipitation followed by sequencing (ChIP-seq) in ES cells ectopically expressing TET3. Binding was widespread throughout the genome and predominantly occurred at promoter regions and CpG islands, but was also enriched at active enhancers and transcribed genes (Fig. 3A, B; antibody specificity Fig. S2B).

**Figure 3.**
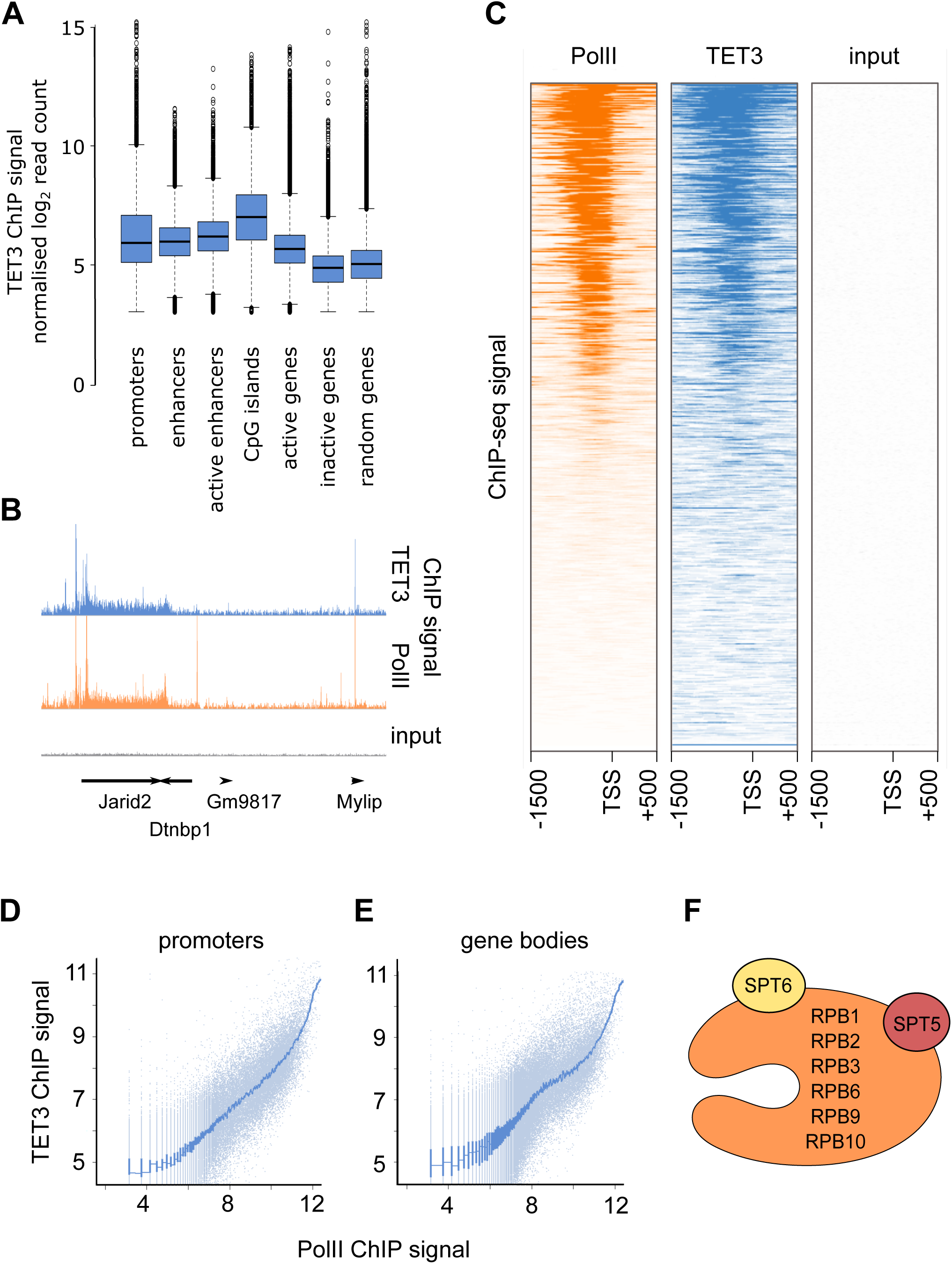
A) Intensity of TET3 ChIP-seq signal over genomic features shown as Tukey box-whisker plots. For clarity the upper limit of the y-axis is set to 15, omitting 31 out of 1,322,843 data points. B) Genome browser shot showing a 700 kb window (GRCm38 chr13:44700000-45400000). Upper and middle tracks show quantification of TET3 ChIP-seq (blue) and PolII ChIP-seq signal (orange), respectively. Signal from material before immunoprecipitation (input) is shown in grey. Location and orientation of genes are indicated by arrows. PolII ChIP-seq data using an antibody which recognises PolII independent of phosphorylation status was taken from (Rahl et al. 2010). C) Aligned probes plot covering a 2 kb region around the transcriptional start site (TSS, -1500 bp to +++500 bp) representing density of ChIP-seq reads by density of colour (orange: PolII ChIP-seq, blue: TET3 ChIP-seq). Each horizontal line corresponds to one genomic region. Lines are sorted by PolII signal intensity with highly transcribed genes featuring at the top of the plot and non-transcribed genes in the bottom half of the plot. ChIP input material does not show any enrichment and is shown in grey. D and E) Quantitative relationship between PolII ChIP-seq signal and TET3 ChIP-seq signal at promoters shown as scatter plot (normalised log_2_ read count). D) Each dot represents a 500 bp window upstream of the transcriptional start site of an mRNA producing gene. E) Each dot represents one gene which shows at least a minimal RNA-seq signal (see Methods: RNA-seq). The blue line depicts smoothed data (median of 250 data points). F) Rough schematic of PolII subunits co-purified with TET3 using rapid immunoprecipitation mass spectrometry of endogenous proteins (RIME, (Mohammed et al. 2013)). RPB1, 2, 3, 6, 9 and 10 are subunits of the core PolII complex, SPT5 and SPT6 are associated factors functioning in transcriptional elongation and mRNA processing. These factors were not present in the IgG negative control (see also Table S2).

Intriguingly, we noted that TET3 was enriched at genomic features known to be bound by RNA Polymerase II (PolII; Fig. EV3; PolII ChIP data from Rahl et al. 2010). Furthermore, the general binding pattern on the genic as well as the genomic level was remarkably similar (Fig. 3B). We therefore explored whether there was a quantitative relationship between TET3 and PolII binding. Indeed, promoters that were not bound by PolII also did not show TET3 occupancy, while progressively higher PolII binding was associated with higher TET3 occupancy (Fig. 3C, D). Of note, TET3 and PolII not only co-occupied genomic regions highly bound by the respective proteins, but also regions which showed moderate PolII enrichment, such as gene bodies of actively transcribed genes (Fig. 3B, E). Co-occupancy of genomic regions by TET3 and PolII was confirmed by rapid immunoprecipitation mass spectrometry of endogenous proteins (RIME, (Mohammed et al. 2013, 2016)) in which immunoprecipitation of TET3 co-purified several subunits of the PolII complex (Fig. 3F, Table S2, antibody specificity Supp. Fig. S2C).

### TET3 enhances chromatin accessibility in RNA Polymerase II bound regions

Prompted by the global increase in transcription and RNA levels, we investigated chromatin accessibility by performing ATAC-seq. This assay exploits the preferential activity of transposase on exposed stretches of DNA in chromatin (Buenrostro et al. 2013). Overall, the pattern of accessible chromatin was very similar between TET3 overexpressing and control ES cells (Fig. 4A, Fig. EV4), which is in line with the observation that the transcriptional pattern does not change dramatically between the samples. Importantly, however, accessible chromatin tended to have a higher ATAC-seq signal in TET3 overexpressing cells (Fig. 4A). Accordingly, for the vast majority of accessible regions identified by MACS peak calling (Zhang et al. 2008) in both sample groups, the ATAC-seq signal was higher in TET3 overexpressing cells (Fig. 4B). When MACS peaks were called independently for control and TET3 overexpressing cells, 85 % of control peaks were found in both samples (8893 out of 10451, Fig. 4C), with an additional 9920 peak regions called in TET3 overexpressing cells. Upon closer inspection, the latter were regions of moderate chromatin accessibility which, while present, did not pass the peak threshold in control cells (Fig. EV4). Moreover, chromatin was significantly more accessible in TET3 overexpressing cells at regions of high PolII occupancy (Fig. 4D), where chromatin is open for transcriptional activity (Fig. 4A) which underlines the link between TET3 and PolII in modulating global transcription. Taken together, ChIP and ATAC-seq results indicate that TET3 colocalises with PolII at open chromatin and enhances accessibility of those regions.

**Figure 4.**
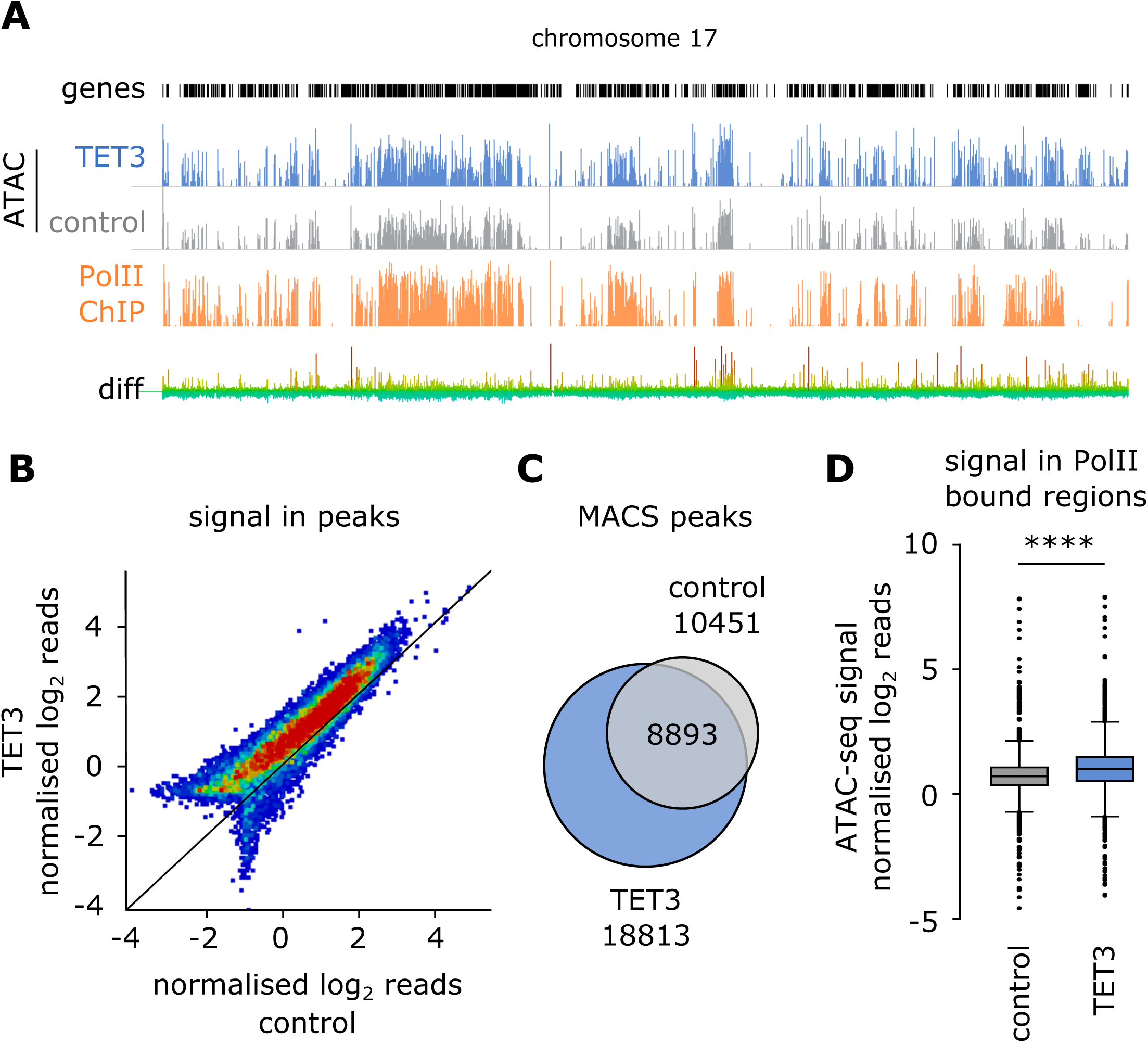
A) Whole chromosome view of chromosome 17. Locations of genes are indicated by black bars. ATAC-seq data for TET3 expressing (top track, blue, y-axis range 0-3) and control ES cells (middle track, grey, y-axis range 0-3), as well as RNA PolII ChIP-seq data (third track, orange, from (Rahl et al. 2010), y-axis range 0-6) quantified as log_2_ read count per million reads over 1 kb running windows. Bottom track shows difference in ATAC-seq read count between TET3 expressing and control cells with gradient colours (green to red) reflecting magnitude of change. B) Density scatter plot showing ATAC-seq reads quantified in all MACS peak regions (control and TET3 overexpressing cells, p<0.00001). For clarity, axes have been scaled and 6 data points out of 20355 are not within the range shown. C) Venn diagram showing numbers of accessibility peaks identified by calling MACS peaks (p<0.00001) independently for TET3 expressing and control cells. D) Tukey box-whisker plots showing ATAC-seq signal in regions bound by PolII (log_2_ read count per million reads > 3 for PolII ChIP-seq) for control (grey) and TET3 expressing (orange) cells. **** p<0.0001, Mann-Whitney test.

### The global effect of TET3 on transcription is independent of catalytic activity and CXXC domain but encoded by evolutionarily conserved exon 4

TET3 is best known for its ability to convert 5-methylcytosine (5mC) to 5-hydroxymethylcytosine (5hmC) and its role in DNA demethylation (Pastor et al. 2013). We therefore investigated whether TET3’s ability to globally enhance transcription was linked to DNA cytosine methylation or its oxidation products. Surprisingly, forced expression of TET3 in ES cells did not result in global changes of 5mC or 5hmC levels (Fig. 5A), which is likely due to the naturally high abundance of TET1 and TET2 in these cells (Yue et al. 2014). To further explore the relationship with DNA methylation we overexpressed a deletion mutant lacking the entire catalytic domain (TET3_trunc_, Fig. 5B, Table S3). Remarkably, this caused the same global shift in exon/intron ratios as full-length Tet3 (Fig. 5C). In contrast, the catalytic domain on its own did not induce the characteristic transcriptional changes (Fig. EV5A). Moreover, like TET3, TET3_trunc_ was also able to enhance conversion of ES to TS-like cells in a transdifferentiation assay (Fig. EV5B) suggesting that catalytic activity was not required to facilitate cellular transitions. Additionally, to rule out the possibility that catalytic activity was recruited through interaction with other TET proteins, we introduced TET3_trunc_ into ES cells deleted for all three TET proteins (TET triple knock out cells, TET TKO, Dai et al. 2016). Analysis of total nascent RNA showed a global increase of transcription in these cells (Fig. 5D). We conclude that the observed effects uncover a novel function of TET3 which is independent of the protein’s catalytic activity or domain.

**Figure 5.**
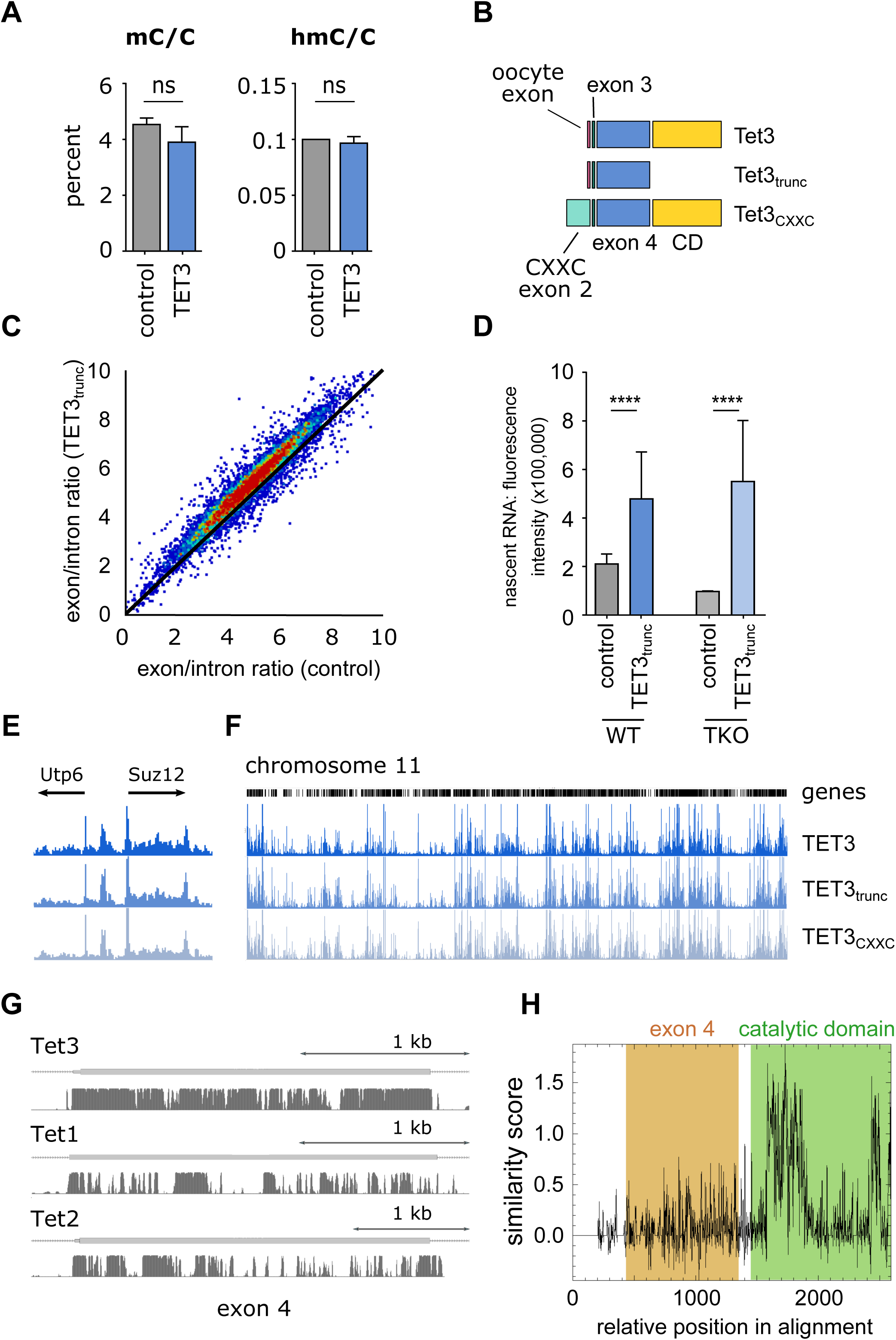
A) Levels of 5-methylcytosine (mC) and 5-hydroxymethylcytosine (hmC) were measured by mass spectrometry in TET3 expressing and control cells and are shown as percentage of total cytosines (C). Bars represent averages with standard deviation (n=3). ns: not significant in a Mann-Whitney test. B) Schematic of Tet3 variants: The oocyte variant consists of an oocyte specific exon (dark red), exon 3 (dark green), exon 4 (blue) and the catalytic domain (CD, yellow, exons 5-11). Exon 1 is non-coding. Tet3_trunc_ is equivalent to oocyte Tet3 without the catalytic domain. Tet3_CXXC_ lacks the oocyte specific exon but contains a CXXC domain encoded in exon 2 (turquoise). Tet3_CXXC_ corresponds to Ensembl transcript Tet3-003 (ENSMUST00000186548.6) and the exon annotation is taken from there. Ensembl Tet3 exon 4 is sometimes referred to as exon 3 or 2 in earlier publications. C) Density scatter plot showing ratios of exonic over intronic reads obtained from total RNA-seq for genes with at least minimum expression (see methods). D) Quantification of fluorescent labelling of nascent RNA in WT and TET triple knock-out cells (TKO) following transient transfection of Tet3. Nascent RNA signal per cell was quantified in TET3 positive (n=95 (WT) and n=139 (TKO)) and control cells (n=115 (wt) and n=29 (TKO)). **** p<0.0001, Mann-Whitney test. E) ChIP-seq signal for TET3 variants in a 128.5 kbp region (GRCm38 chr11:79925225-80053751). Top row represents oocyte TET3, second row truncated TET3 and last row somatic TET3 containing the CXXC domain. Genes and direction of transcription are indicated by arrows. Data were quantified as per million normalised read count per 1 kb window. Y-axis range is 0-3. F) ChIP-seq profiles for TET3 variants on chromosome 11. Genes are indicated by black bars. Data were quantified as log_2_ read count per 1 kb window normalised per million reads. Y-axis range is 0-3. Top row represents oocyte TET3, second row truncated TET3 and last row somatic TET3 containing the CXXC domain. G) Evolutionary conservation across placental mammals of Tet3 exon 4 and corresponding exons of other Tet family members was measured using the PhastCons (Siepel et al. 2005). Orange bars represent exons with the scale indicated. Conservation scores between 0 and 1 are represented by the height of the black bars. H) Amino acid similarity across proteins of the TET family visualised using EMBOSS:plotcon (Rice et al. 2000). Amino acids of the catalytic domain are highlighted in green, amino acids encoded in exon 4 harbouring a transcriptional activation function of TET3 are highlighted in orange. The region encoded in exon 4 is less similar between TET proteins than the catalytic domain.

TET3 has previously been shown to partner with O-linked N-acetylglucosamine transferase (OGT) thereby influencing transcription independent of its oxidising function (Vella et al. 2013; Chen et al. 2013; Deplus et al. 2013; Ito et al. 2014). We confirmed TET3’s strong interaction with OGT by RIME (Table S2), however, this interaction was lost upon deletion of the catalytic domain (Fig. EV5D, Table S4, Deplus et al. 2013; Ito et al. 2014). Furthermore, genes identified by RNA-seq as upregulated were still activated by TET3 upon OGT knock-down (Fig. EV5E). Thus, the novel function described here is not mediated by the interaction of TET3 and OGT.

We next aimed to determine if the TET3 variant produced in oocytes was functionally different from the somatic variant. The most striking difference between the protein variants is the presence or absence of a CXXC domain encoded in exon 2 (Fig. 1A, 5B, Table S3). While in somatic tissues the CXXC domain is predominantly present (referred to as TET3_CXXC_), this exon is skipped in the Tet3 transcript found in oocytes (referred to as TET3). CXXC domains bind CpG rich sequences in DNA (Long et al. 2013) and have thus been proposed to be responsible for TET protein targeting. We therefore refer to the somatic variant as TET3_CXXC_. Strikingly, when we compared genomic binding profiles of TET3 and TET3_CXXC_ by ChIP-seq, we found them to be virtually identical (Fig. 5E, F, EV5C) arguing that the CXXC domain does not influence targeting of TET3 in this system. Moreover, forced expression of TET3_CXXC_ induced the same transcriptional changes as TET3 (Fig. EV5F) for a panel of genes affected by TET3 overexpression.

We attempted to further narrow down the region responsible for TET3’s transcriptional function. Tet3_trunc_ consisting of the oocyte exon, and exons 3 and 4 (Fig. 5B) was further truncated to exon 4, as the other two encode only 11 and 19 amino acids, respectively. Indeed, specific overexpression of exon 4 alone was able to elicit the same transcriptional effect on genes affected by overexpressing full-length Tet3 (Fig. EV5G). Interestingly, while generally exons from Tet genes are similarly conserved across placental mammals, exon 4 displays much higher conservation scores than the corresponding exons in Tet1 and Tet2 (Fig. 5G, EV5H). Furthermore, comparison of TET1/2/3 at the protein level revealed that while the amino acid sequences of their catalytic domains are very similar, the sequences encoded by the large exon 4 are not (Fig. 5H). Overall, this supports the idea that Tet3 exon 4 carries a conserved function that is unique within the TET proteins.

### Discussion

Defining and preserving cell identity is crucial for multicellular organisms. Once established, cell identity is protected by several levels of epigenetic regulation including DNA methylation, chromatin modifications and remodelling, and spatial arrangement of the genome (Barrero et al. 2010; Allis and Jenuwein 2016). However, cell identity must not be immutable as the life cycle of an organism requires cells to transition between states. Often these cell fate transitions entail changes in developmental potency of the cell and are accompanied by major alteration of the chromatin environment and transcriptional landscape (Chen and Dent 2014; Lee et al. 2014; Apostolou and Hochedlinger 2013). The beginning of a new generation marks the most extreme case of cell identity change: Upon fusion of two highly specialised, transcriptionally quiescent cells, the totipotent zygote is created which completely re-organises the parental genomes to start its own transcriptional program (Clift and Schuh 2013; Borsos and Torres-Padilla 2016; Percharde et al. 2017a). Much more frequent than this unique event, however, are cell identity changes during differentiation that restrict cellular potency. These occur for example at the exit from pluri- or multipotency during development and adult life of an organism (Krishnakumar and Blelloch 2013; Lee et al. 2014; Soufi and Dalton 2016). A common theme to all these transitions is widespread restructuring of chromatin and the rewiring of transcriptional networks. It has recently been proposed that during such transitions, global transcription is temporarily upregulated, a phenomenon termed hypertranscription (Percharde et al. 2017a). Here, we provide evidence that TET3 increases transcription, RNA levels and chromatin accessibility genome-wide and may thereby be involved in promoting changes of cell identity.

In this manuscript we show examples of Tet3 being expressed or upregulated during major cellular transitions. Tet3’s prominent presence in very early embryos can be viewed in this light: Highly abundant Tet3 transcript in oocytes/zygotes is rapidly and completely depleted by the 4-cell stage (Tan and Shi 2012) compatible with a function during this profound cellular reorganisation that later on is no longer required and possibly detrimental. Interestingly, global transcriptional activation mediated by TET3 also resulted in the upregulation of MERVL elements which orchestrate expression of a network of early embryonic genes (Macfarlan et al. 2011) critical for early preimplantation development (Kigami et al. 2003). This is in line with the possibility of TET3 playing a role in zygotic genome activation. Other examples of TET3 upregulation when cells undergo identity changes include exit from pluripotency (Veillard et al. 2014), iPS reprogramming (Milagre et al. 2017), SCNT reprogramming (Hosseini et al. 2016) and PGC specification (Seisenberger et al. 2012). In fact, PGCs have recently been shown to exhibit increased RNA levels, elevated transcription, increased cell size and upregulation of ribosomal protein transcripts (Percharde et al. 2017b), features that are mirrored in Tet3 expressing ES cells. An active role rather than a mere correlation is supported by the ability of ectopically expressed TET3 to enhance transdifferentiation from ES to TS cells, a system in which TET3 is not normally present. Our findings support the proposal that global hypertranscription may be an important component of cell identity changes during development (Percharde et al. 2017a).

Our results group TET3 with other factors globally enhancing transcription. For example, the chromatin remodeller Chd1 has been shown to enhance the activity of PolII by removing nucleosomal barriers (Skene et al. 2014; Guzman-Ayala et al. 2015). Upon deletion, global transcriptional output is reduced and embryos fail to sustain epiblast development (Guzman-Ayala et al. 2015). In a mechanistically different fashion, the global regulator c-Myc amplifies transcription in cancer cells by stimulating transcriptional pause release via P-TEFb (Rahl et al. 2010). Interestingly, hypertranscription in PGCs is also dependent on Myc/Max and P-TEFb (Percharde et al. 2017b). It will be interesting to explore further whether TET3 uses the same pathways or acts in a parallel manner.

In contrast to our expectations, we found that the transcriptional changes brought about by the oocyte and somatic isoforms of Tet3, as well as their chromatin occupancy were almost indistinguishable. This argues against a specific function of TET3 in the oocyte/zygote but supports the idea that TET3 generally promotes cellular transitions. Like many genes, Tet3 has an oocyte specific promoter (Veselovska et al. 2015) which likely functions to ensure temporally restricted high expression of Tet3 rather than producing a protein with a different function.

TET3 is clearly a multifunctional protein and its oxidase activity is only one mode of action. Several other reports have shown additional non-catalytic functionalities of the TET proteins (for an overview see (Lian et al. 2016, Figure 3). Most notably, like TET1 and TET2, TET3 interacts strongly with O-linked N-acteylglucosamine transferase (OGT) and plays an important role in the recruitment of OGT to chromatin, where it influences transcription through GlcNAcylation of histones and chromatin modifying complexes such as SET1/COMPASS (Vella et al. 2013; Chen et al. 2013; Deplus et al. 2013; Ito et al. 2014). However, the transcription and chromatin changes induced by TET3 are independent of its association with OGT. TET3 has also been shown to interact with REST and H3K36 methyltransferases to mediate transcriptional activation (Perera et al. 2015). Additionally, TET3 activity is regulated by CRL4 (Yu et al. 2013) and its intracellular location affected by post-translational modification through OGT (Zhang et al. 2014). It is noteworthy that the transcriptional function of Tet3 is independent of its catalytic and CXXC domains since other previously reported non-catalytic activities of TET proteins are still mediated by the catalytic domain (Deplus et al. 2013; Yu et al. 2013; Zhang et al. 2014; Ito et al. 2014).

Interestingly, and presumably a result of the multiple global activities of TET3, previous studies have reported different and sometimes contradictory loss of function effects. The comparison is potentially confounded by the use of different knock-out strategies which have deleted different parts of the transcript (Table S5). Given the evidence for a functional truncated transcript *in vivo* (ensemble transcript Tet3-002, ENSMUST00000056191.1, HAVANA project), certain knock-out designs may not result in the absence of the entire protein and therefore preserve some of the non-catalytic functions of TET3. For instance, the knock-out generated in our lab produces a frame shift and premature stop codon upstream of the catalytic domain (Santos et al. 2013; Peat et al. 2014), and while this completely abolishes catalytic activity, a truncated transcript including exon 4 is still present and translated. Two separate studies have deleted exon 4 (referred to as exon 3 in (Tsukada et al. 2015) and exon 2 in (Kang et al. 2015)). Interestingly, Tsukada and colleagues report that TET3 contributes to the fine-tuning of zygotic transcription after DNA synthesis, although their findings point to an inhibitory effect. Kang and colleagues report increased transcriptome variability in Tet1/3 double knock-outs in individual 8-cell blastomeres and blastocysts, concomitant with variable delayed or aborted development.

Taken together, our findings support TET3 as a multifaceted protein with potentially complementary functions; whereas the C-terminal half promotes the resetting of epigenetic marks, we provide evidence that the largely uncharacterised exon 4 can globally increase transcription. While *in vitro* temporary hypertranscription is sufficient to facilitate cellular transitions, *in vivo* it may function in concert with the removal of DNA methylation and deposition of chromatin modifications. We propose that TET3’s combined functions create a favourable chromatin environment for transitioning between cell identities. Importantly, we envision TET3 as a facilitator rather than an driver of change which still allows the protein to exert its catalytic function in systems not primed for change, for example the brain (Hahn et al. 2013). It is clear we are only beginning to understand the complex interplay of chromatin remodelling, epigenetic marks on DNA, and global transcription that enable change of the epigenetically well-guarded cellular identity.

### Materials and Methods

#### Cloning of Tet3 variants

Tet3 variants were cloned from cDNA using MII oocytes (oocyte specific Tet3) or embryoid bodies (Tet3_CXXC_) and recombined into pDONR221 (Invitrogen) by Gateway cloning. Several expression constructs were used: Constitutive strong expression of the Tet3 transgene was achieved using vectors based on PB-DST-BSD which was a kind gift from Jose Silva. Transgenes can be inserted into this backbone via Gateway cloning and can then be expressed from a CAG promoter. Resulting plasmids from the pIG300 series (this study) do not contain a fluorescent marker, plasmids from the pIG400 series (this study) create a C-terminal eGFP fusion for the transgene. Inducible Tet3 expression was achieved using constructs based on PB_TAG_PB_tetO2_iresGFP which was a kind gift from Peter Rugg-Gunn (pIG200 series in this study). Transgenes can be inserted via Gateway cloning, and expression is driven by a tetracycline responsive CMV minimal promoter with eGFP being expressed from an internal ribosome entry site (IRES). A list of plasmids used for the generation of cell lines is provided in Table S6 and sequences are available on request.

#### Cell culture and cell line construction

E14 mouse embryonic stem cells were grown under standard serum/LIF conditions (DMEM, 4,500 mg/l glucose, 4 mM L-glutamine, 110 mg/l sodium pyruvate, 15 % fetal bovine serum, 1 U/ml penicillin, 1 mg/ml streptomycin, 0.1 mM nonessential amino acids, 50 mM b-mercaptoethanol, and 1000 U/ml LIF). TET3 KO cells and the parental cell line were a gift from Fabio Spada and Guo-Liang Xu and were cultured under the following conditions: DMEM, 4,500 mg/l glucose, 4 mM L-glutamine, 110 mg/l sodium pyruvate, 20 % fetal bovine serum, 1 U/ml penicillin, 1 mg/ml streptomycin, 0.1 mM nonessential amino acids, 50 mM b-mercaptoethanol, 2 mM GlutaMAX (Gibco), 1000 U/ml LIF, 1 μM PD0325901 and 3 μM CHIR99021. E14 ES lines harbouring Tet3 variants were generated by transfection with FuGENE6 (Promega), plasmid integration via the piggyBAC system (Kim and Pyykko 2011) and selection with the relevant drug. Control cell lines carry the empty vector and were equally selected. For the inducible expression system, piggyBAC vectors were co-transfected with a plasmid carrying a reverse tetracycline responsive transactivator (rtTA2s-M2). Unless otherwise indicated the pool of stable integrants was used, and TET3 expressing cells enriched by flow sorting for GFP fluorescence using a BC Influx High-Speed Cell Sorter. For the generation of clonal lines individual colonies were picked and expanded.

#### RNA isolation, qPCR and total RNA-sequencing

RNA was isolated using Qiagen RNeasy Micro columns and treated with DNaseI (Ambion). cDNA was generated using 0.5 – 1 μg RNA (Invitrogen SuperScript II) and qPCR performed using the Brilliant III SYBR mix (Agilent Technologies). Relative quantification was performed using the comparative CT method with normalisation to Hsp90 and Atp5b levels. Primer sequences are available on request. Opposite strand-specific total RNA libraries with RiboZero rRNA depletion (Illumina TruSeq Kit) were prepared from 200 ng – 1 μg DNase treated RNA by the Sanger Institute Illumina bespoke pipeline. Sequencing was performed as paired-end 75 bp runs on the Illumina HiSeq 2500 Rapid Run platform. At least three biological replicates were used, generating 28-177 × 10^6^ mapped reads per sample (Table S7).

#### hmC and mC mass spectrometry

Approximately 350 ng DNA was digested to single nucleotides using the DNA Degradase Plus kit (Zymo). LC-MS/MS was performed on a Q-Exactive mass spectrometer (Thermo Scientific) fitted with a nanoelectrospray ion-source (Proxeon). Mass spectral data were acquired in MS/MS mode at a relative collision energy of 10%, selecting the parent ions at m/z 228.1 (C), 242.1 (mC) and 258.1 (hmC) with a 1 amu isolation width. Fragment ion spectra were acquired over the m/z range 100-300 at a nominal resolution setting of 35,000 (at m/z 200), and peak areas for the fragment ions 112.0505 (C), 126.0662 (mC) and 142.0611 (hmC) were obtained from extracted ion chromatograms of the relevant scans.

#### RNA-seq analysis

Total RNA-Seq reads were trimmed using Trim Galore v0.4.2 (http://www.bioinformatics.babraham.ac.uk/projects/trim_galore/) using default parameters to remove the standard Illumina adapter sequence. They were then mapped to the mouse GRCm38 genome assembly using Tophat 2.0.12 guided by the gene models from the Ensembl v70 release.

Figure 2: Prior to RNA isolation cell lines carrying different Tet3 constructs were enriched for cells with TET3 expression by flow sorting for GFP fluorescence using a BD Influx High-speed Cell Sorter. Data analysis was performed in SeqMonk (http://www.bioinformatics.babraham.ac.uk/projects/seqmonk/) using the following parameters: Reads in exons were quantitated with the RNA-seq quantitation pipeline, counts normalised per million total reads per sample and log_2_ transformed. Raw counts were used for DESeq2 analysis (Love et al. 2014). Differentially expressed genes were identified using the intersection of DESeq2 statistics (FDR<0.05) and an intensity difference filter (p<0.05 with Benjamini and Hochberg multiple testing correction, https://www.bioinformatics.babraham.ac.uk/projects/seqmonk/Help/5%20Filtering/5.2%20Statistical%20Filters/5.2.4%20Intensity%20Difference%20Filter.html). Functional enrichment analysis was performed using g:profiler (http://biit.cs.ut.ee/gprofiler/).

Figure S1: Differentially expressed genes were identified using the intersection of DESeq2 statistics (FDR<0.05) and a dynamic fold-change filter (https://www.bioinformatics.babraham.ac.uk/projects/seqmonk/Help/5%20Filtering/5.2%20Statistical%20Filters/5.2.4%20Intensity%20Difference%20Filter.html). For the comparison of groups ‘up’, ‘down’ and ‘random’ only genes were used which showed a minimum coverage by RNA-seq reads (log_2_ read count > -2). For quantitative data (e. g. gene length) two random sets of genes were generated of which one is shown (random samples were very similar throughout). For categorical data (e. g. fraction of paused genes), three random sets of genes were generated and average with standard deviation is shown. Statistical analysis was performed in Seqmonk or GraphPad Prism 6.

#### ChIP-Seq

Chromatin immunoprecipitation was performed essentially as described in (Schmidt et al. 2009). 6 15 cm dishes of ES cell lines expressing different TET3 isoforms (301, 305, 307, see Table S6) were cross-linked and harvested. 10 ug of a TET3 antibody developed with Millipore (ABE290) was used per precipitation, and cross-linked, sonicated nuclear material was used as input control. Libraries for Illumina sequencing from precipitated DNA were prepared using the Diagenode MicroPlex Library Preparation Kit and were sequenced on 2 lanes of an Illumina HiSeq2500 as 50 bp single-end reads. For samples and their respective read counts see Table S7.

ChIP-Seq reads were trimmed using Trim Galore v0.4.2 (http://www.bioinformatics.babraham.ac.uk/projects/trim_galore/) using default parameters to remove the standard Illumina adapter sequence. They were mapped to the mouse GRCm38 genome assembly using Bowtie 2 (v2.2.5, default parameters).

Figure3: Data analysis was performed in SeqMonk (http://www.bioinformatics.babraham.ac.uk/projects/seqmonk/) using the following parameters: Reads were quantitated in running windows (1 kb), counts normalised to the total number of sequences per sample and log_2_ transformed. Probes were filtered for those showing at least some background binding (normalised log_2_ read count per 1 kb >3). ChIP-seq data for N-terminal RNA PolII was taken from (Rahl et al. 2010). Feature analysis was performed using the following annotations: promoters: 500 bp upstream of mRNA; enhancers: H3K4me1 signature from (Creyghton et al. 2010); active enhancers: H3K27ac signature from (Creyghton et al. 2010); active genes: RNA PolII bound promoters (data from (Rahl et al. 2010) and transcribed (RNA-seq data from this study); inactive genes: non bound by RNA PolII (data from Rahl et al., 2010) and not transcribed (RNA-seq data from this study). Statistical analysis was performed in Seqmonk or GraphPad Prism 6.

#### ATAC-seq

ATAC-seq was performed as per (Buenrostro et al. 2013) using 10,000 cells per each of three biological replicates and 15 cycles of PCR amplification. Samples were barcoded, pooled and sequenced across two lanes of a HiSeq2500 as 75bp paired-end reads. For samples and their read counts see Table S7. ATAC-seq reads were quantified as log_2_ read count per million reads over 1 kb running windows. Visualisation, quantitation and statistical analysis was performed in Seqmonk and GraphPad Prism 6.

#### Conservation analysis

Conservation scores across placental mammals for Tet family members were obtained using the PhastCons track in the UCSC genome browser (Siepel et al. 2005) and a zoomed in view of Tet3 exon 4 and corresponding exons of other Tet genes is shown in Fig 4. Similarity at protein levels between TET proteins was analysed using plotcon (http://emboss.sourceforge.net/apps/release/6.6/emboss/apps/plotcon.html).

#### Nascent transcription imaging assay

Newly synthesised RNA was visualised using the Click-iT Plus Alexa Fluor Picolyl Axide Toolkit (Molecular Probes) according to instructions of the manufacturer. In brief, cells were pulsed with 5-ethynyl uridine (EU) at 1 mM for 20 minutes and then fixed in 2 % formaldeyde for 30 min at room temperature. Cells were cytospun onto poly-lysine slides and permeabilised using PBS/0.5% triton X-100 for 30 min. TET3 was visualised by immunofluorescence using anti-TET3 antibody (ab290, Millipore, 1:500) as primary and anti-rabbit Alexa Fluor 647 (A31573, Molecular Probes, 1:500) as secondary antibody. EU incorporated into RNA was labelled with picolyl azide Alexa Fluor 555 via Click-iT reaction using a 1:1 ratio of CuSO4 and copper protectant. DNA was stained using DAPI (1:1000). Imaging was done on a Nikon A1R+ resonant scanning confocal microscope and signal intensities were quantified using Volocity software. To exclude effects due to slide to slide variation of signal intensity, TET3 expressing and control cells were mixed on slides at the beginning of the experiment and later identified by TET3 immunofluorescence. 5EU pulsing was performed twice and up to five slides were stained per experiment. Absolute fluorescence was different between slides (as expected) but overall results are consistent. Statistical analysis was done with GraphPad Prism 6 and signal distributions on one slide are shown.

#### Quantification of RNA content

Defined numbers of TET3 expressing and control cells were collected by flow cytometry using a BD Influx High-speed Cell Sorter. Five independent sorts were performed on cells from different passages and the same number of cells collected for both cell lines. Numbers per experiment were 116, 200, 200, 180 and 181 × 10^3^. For each cell line only cells showing high GFP fluorescence were collected with gates constant between cell lines and experiments. RNA was isolated using Qiagen RNeasy Micro Kit according to the instructions of the manufacturer (which includes DNase treatment). RNA was quantitated as 1:100 and 1:200 dilutions using the Qubit RNA HS Assay Kit (Thermo Fisher Scientific).

#### Transdifferentiation assay

TS base media consisting of RPMI 1640 (Gibco) supplemented with 20 % FBS (Gibco), 1 mM sodium pyruvate (Gibco), 50 U/ml penicillin-streptomycin (Gibco) and 0.05 mM beta-mercaptoethanol (Gibco) was conditioned by incubation with irradiated MEF cells on cell culture dishes for two days and then passed through a 0.22 μm filter. Complete TS cell medium was prepared by combining 70 % conditioned media, 30 % TS base media, 20 ng/ml beta-foetal growth factor and 1 μg/ml heparin. The indicated ES cell lines were seeded at a density of 400 cells/cm^2^ onto a layer of irradiated MEF cells in full TS medium without antibiotic selection. Media was changed every 48 hours. After six days, cells were first visually examined by phase-contrast microscopy and then harvested with trypsin. Cells were incubated with anti-CD40 antibody (AF440, R&D systems, 1:50) for 30 min, then washed and incubated with anti-goat Alexa Fluor 647 antibody (A21447, Thermo Fisher Scientific, 1:500) and anti-THY1-PE (12-0900-81, eBioscience, 1:2500, feeder detection). To assess cell viability, 1 μg/ml DAPI (Sigma) was added. Immunofluorescent signal was then analysed on a BD LSRFortessa flow sorter.

#### RIME

RIME proteomics experiments were performed as previously described (Mohammed et al. 2013, 2016). For SILAC RIME experiments E14 ES cells were grown in R/K deficient SILAC DMEM (PAA E15-086) and supplemented with 800 μM L-Lysine ^13^C_6_^15^N_2_ hydrochloride and 482 μM L-Arginine ^13^C_6_^15^Nhydrochloride (Sigma-Aldrich) for “heavy”-labelled media or 800 μM L-Lysine ^13^C_6_^15^N_2_ hydrochloride and 482 μM L-Arginine ^12^C_6_^14^N_4_ hydrochloride for “light”-labelled media. Table S2 contains metrics for proteins that were identified by proteomics after TET3 pull-down but not in the IgG control. Table S4 contains proteins identified by SILAC RIME and the log_2_ ratio of peptide counts of TET3 over TET3_trunc_ samples.

## Acknowledgements

We thank Ines Milagre, Lenka Veselovska and Sebastien Smallwood for sharing RNA-seq data before publication. We would also like to thank Francesco Cambuli, Dominika Dudzinska and Myriam Hemberger for reagents and advice on the transdifferentiation assays, Felix Krueger for bioinformatics processing, David Oxley for mass spectrometry analysis of nucleosides, Hanneke Okkenhaug for quantification of fluorescent imaging signals and Anne Segons-Pichon for statistical advice. We thank Millipore for the generation of the TET3 antibody, and Fatima Santos for initial tests. We thank the Wellcome Trust Sanger Institute Bespoke Sequencing Team for RNA-seq library preparation and sequencing, the Proteomics Core Facility at the Cancer Research UK Cambridge Institute and the Babraham Institute Flow Cytometry Facility. We thank Fabio Spada and Guo-Liang Xu for TET3 TKO cells. We are grateful to Irene Min and John Lis for sharing their lists of genes for RNA Polymerase pausing categories. We thank all members of the Reik lab for helpful discussions. This work was funded by the Wellcome Trust (095645/Z/11/Z) and the BBSRC (BB/K010867/1). ME-M is supported by a Marie Sklodowska-Curie Individual Fellowship and JP by the Rutherford Foundation Trust.

## Accession numbers

A GEO reviewer link has been created for record GSE94688.

## Author contributions

CK, JP and WR conceived and designed the study. CK performed experiments, analysed data and wrote the paper. JP performed experiments and analysed data. ME-M performed experiments, analysed data and edited the paper. TH and JP characterised the oocyte isoform of Tet3 and did evolutionary comparisons. HM performed RIME experiments. SA helped analyse sequencing data. WD and WR helped interpret data and supervised the study. WR edited the paper.

## Expanded View Figure Legends

**Figure EV1.**
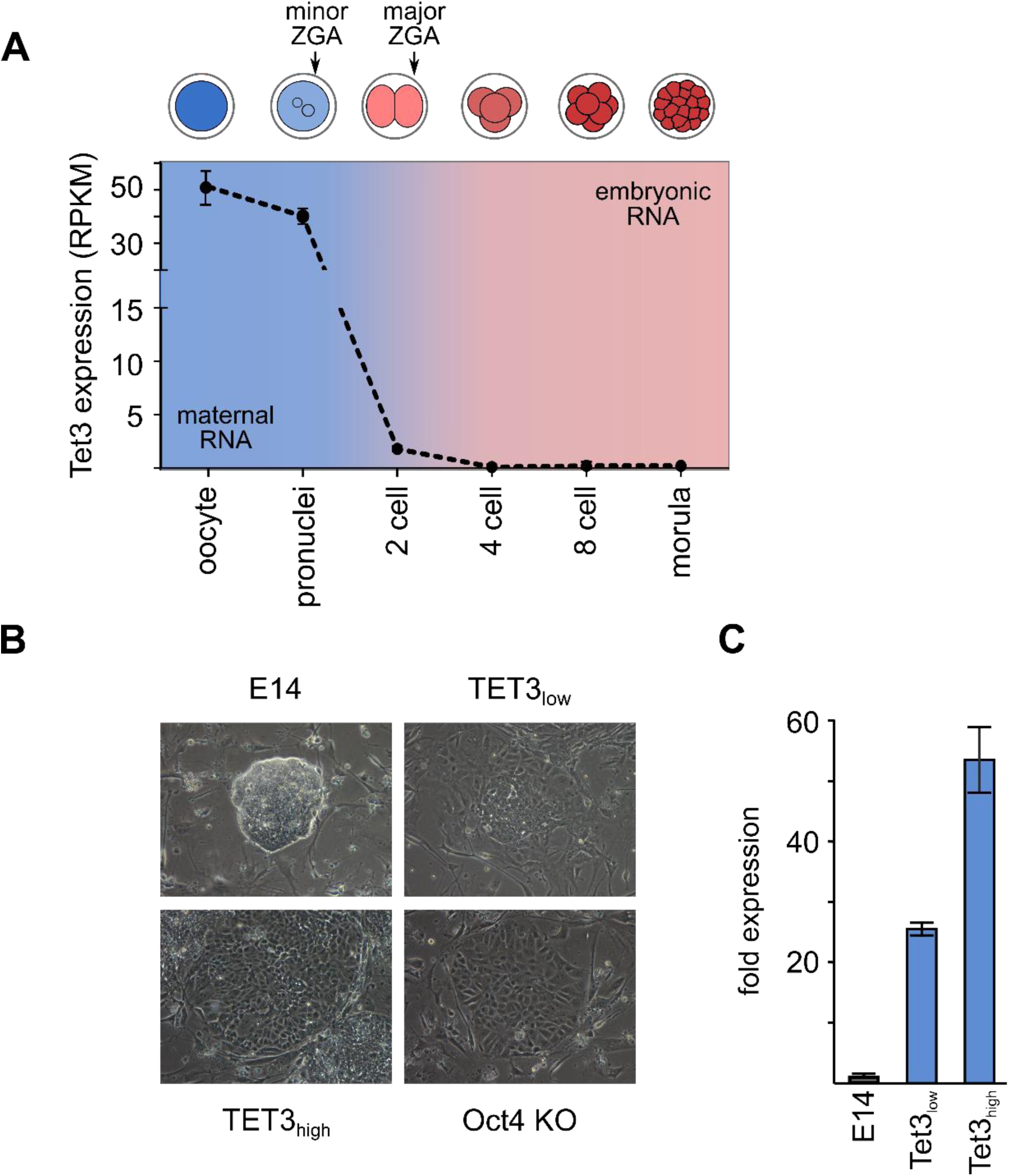
A) Tet3 expression during early embryonic development monitored by RNA-seq (RPKM: reads per kilobase per million mapped reads; data from Xue et al. 2013). Tet3 RNA is provided by the oocyte and is rapidly degraded after fertilisation. Stages of cleavage embryos are schematically shown above the graph. Activation of the zygotic genome (ZGA) happens in a minor and a major wave in late zygotes and 2-cell embryos, respectively. The bulk of stored maternal RNA is degraded by the 2-cell stage. B) Example images of cells after six days of trans-differentiation. Some of the TET3 positive colonies display a flat, epithelial like morphology. Oct4 knock-out ES cells (Oct4 KO, ZHBTc4 cells, (Niwa et al. 2000, 2005)) are a model for facilitated trans-differentiation. TET3_low_ and TET3_high_ denote clonal cell lines which express different amounts of TET3. C) Tet3 expression levels determined by qPCR for cell lines used in transdifferentiation experiments. Control cells are E14 mouse ES cells, TET3_low_ and TET3_high_ are clonal lines derived from E14 cells stably transfected with Tet3 (see Table S6). Bars represent the mean of technical duplicates with error bars indicating the range. Tet3 expression in E14 was set to 1. Differential TET3 expression was confirmed visually by immunofluorescence.

**Figure EV2.**
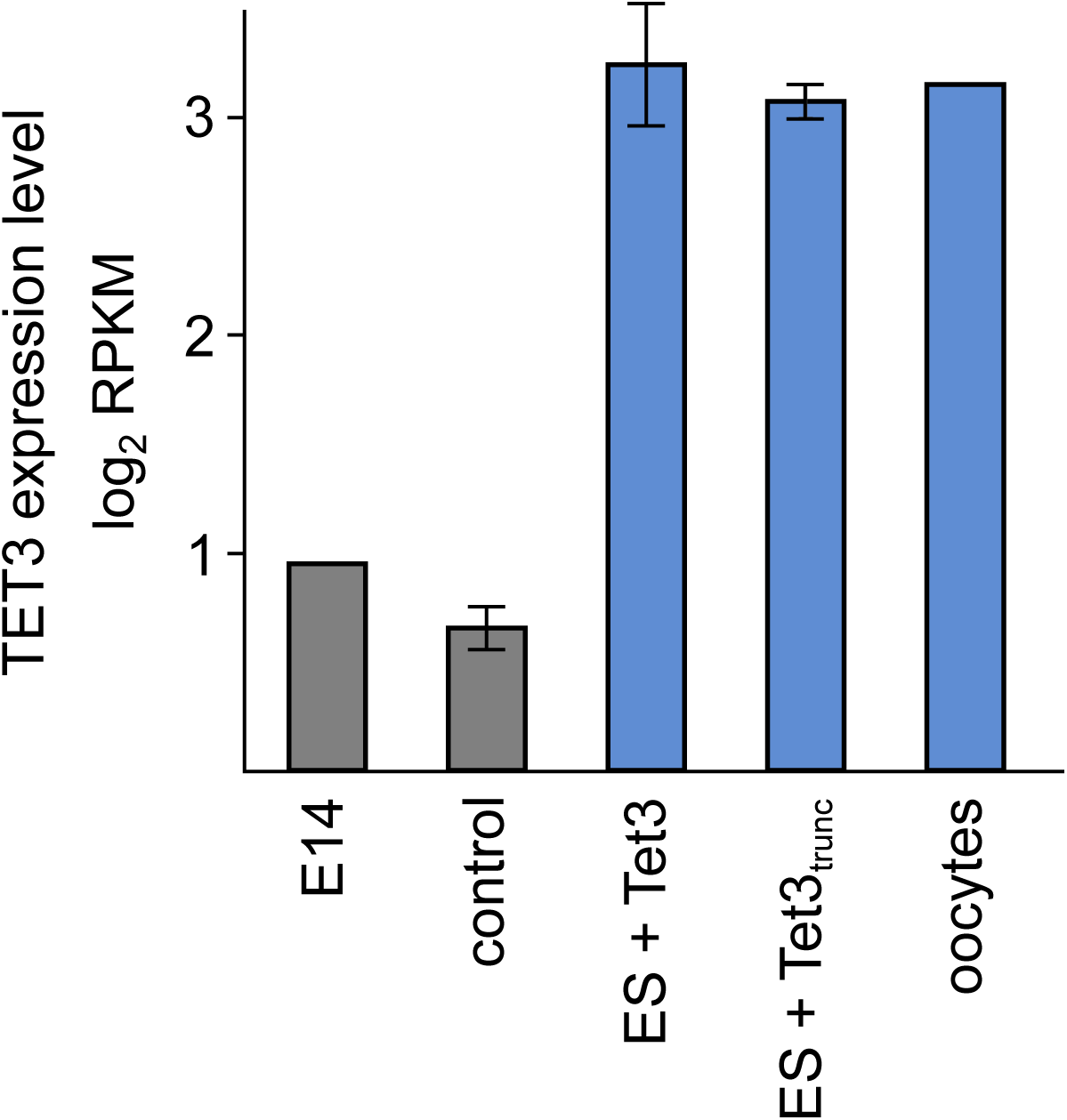
Comparison of Tet3 expression levels in different cell lines and oocytes quantified by RNA-seq. Reads were counted over the region common to all variants. E14 are WT ES cells. Control cells are E14 transfected with the empty vector. ES + Tet3 and ES + Tet3_trunc_ are stable cell lines constitutively expressing Tet3 variants (Table S6). Oocyte expression data is from (Veselovska et al. 2015; FGO). Bars represent mean log_2_ RPKM, error bars indicate standard deviation for samples in which three biological replicates were available.

**Figure EV3.**
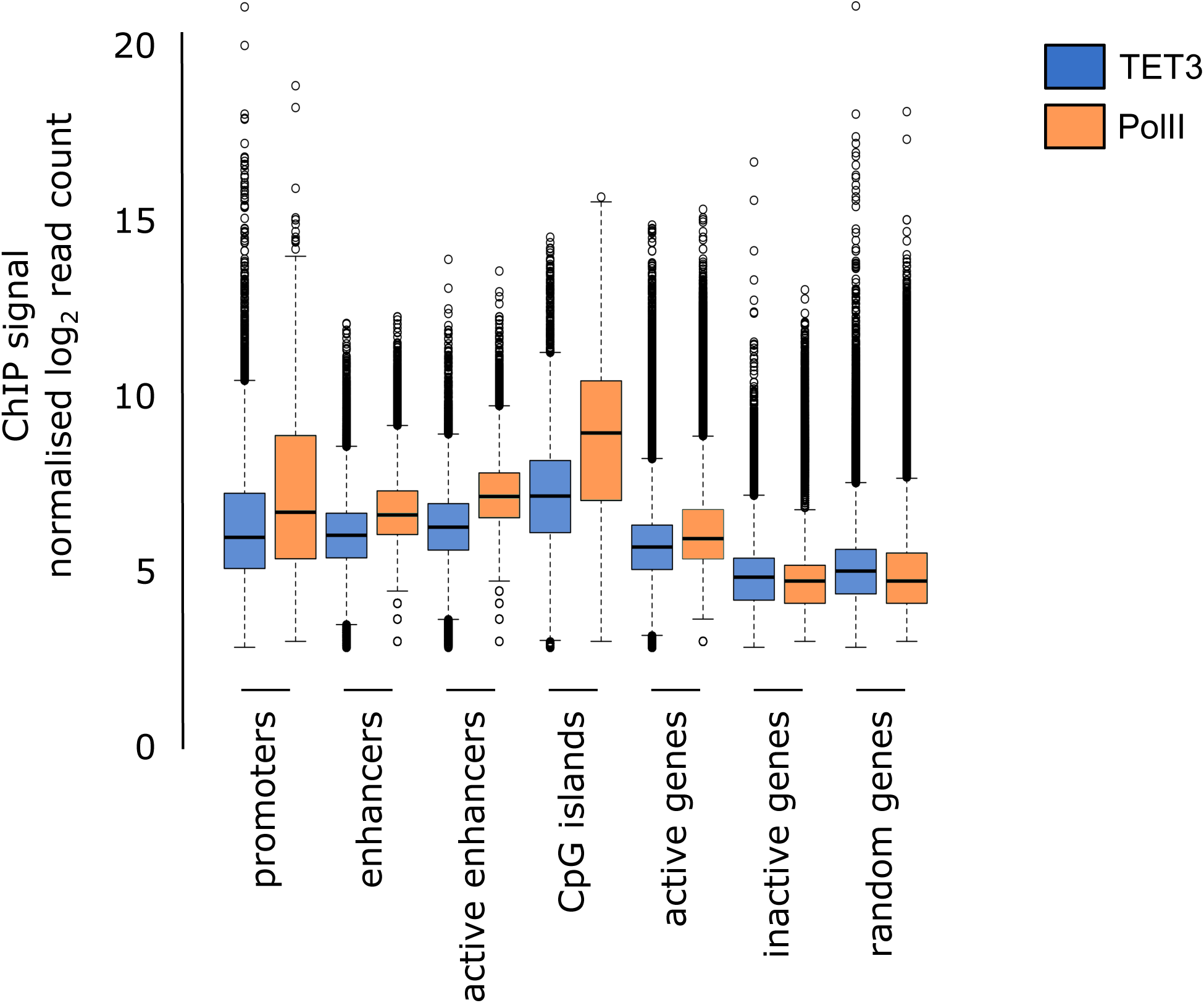
Intensity of TET3 (blue) and RNA PolII (orange) ChIP-seq signal over genomic features shown as Tukey box-whisker plots. Data for RNA PolII binding was taken from (Rahl et al., 2010).

**Figure EV4.**
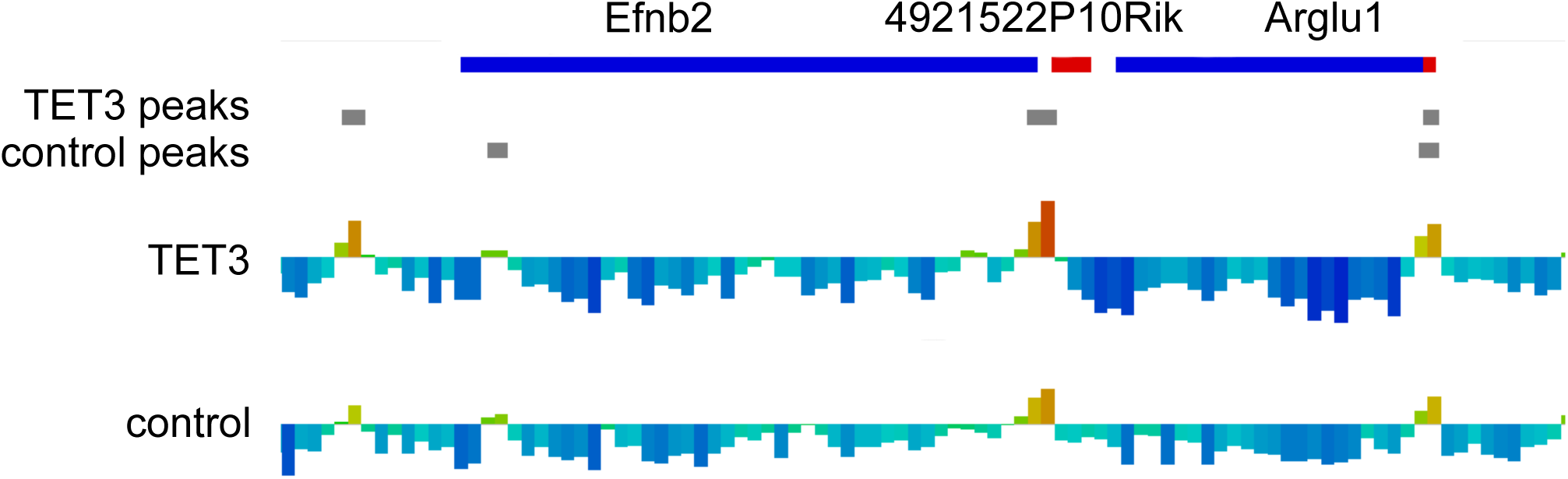
Genome browser shot of ATAC-seq data quantified as log_2_ read count per 1 kb window normalised per million reads for region GRCm38 chr8:8604116-8700350. Quantification is visualised by heat colours (blue: low, red: high) and height of the bar. MACS peaks were called for TET3 overexpressing and control cells and are indicated as grey, horizontal bars. Sense and antisense genes are drawn as red and blue boxes, respectively. Note that even when a peak is called for the TET3 expressing but not the control sample, ATAC-seq reads are still enriched in this region arguing for a quantitative rather than a qualitative difference between samples.

**Figure EV5.**
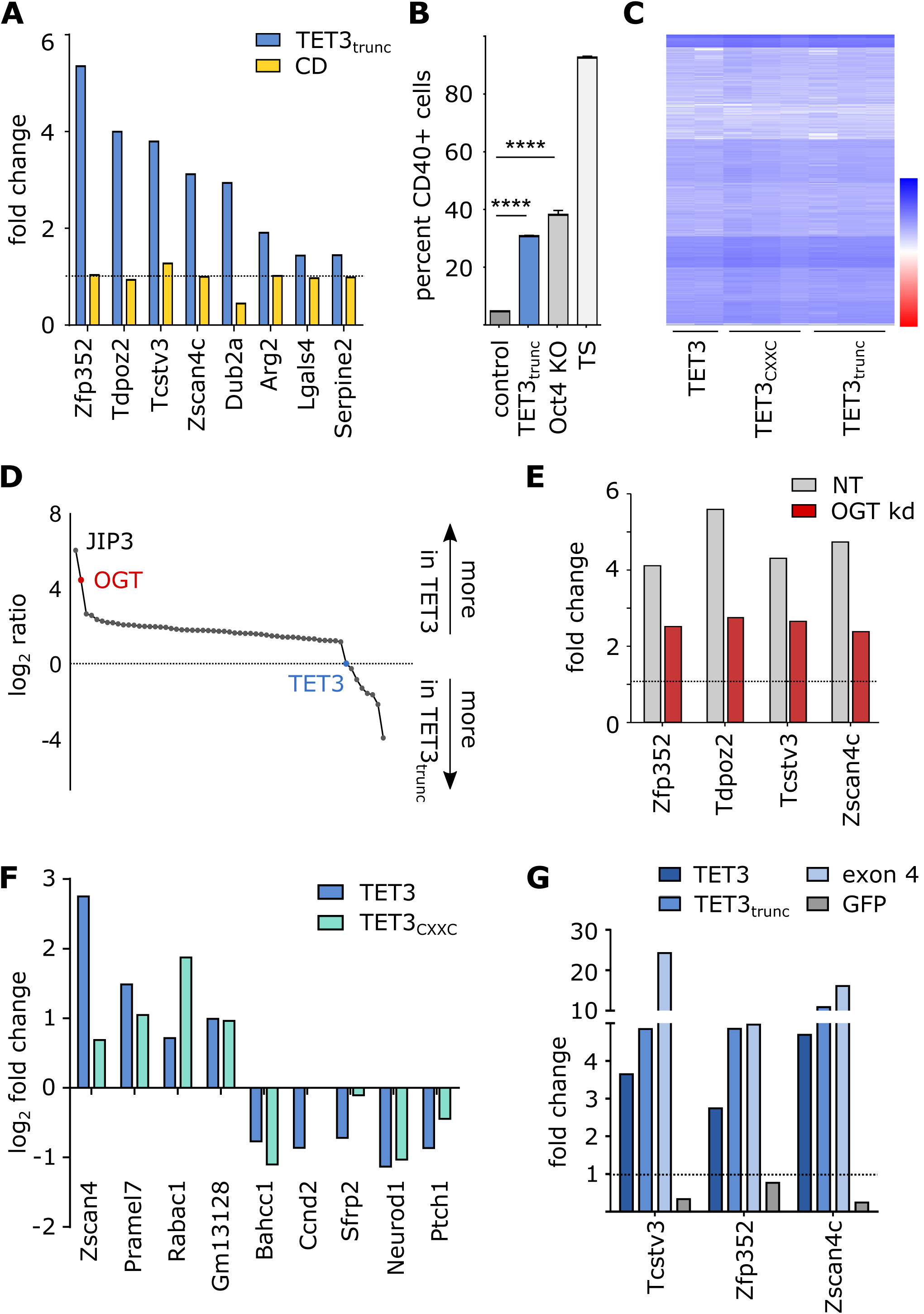
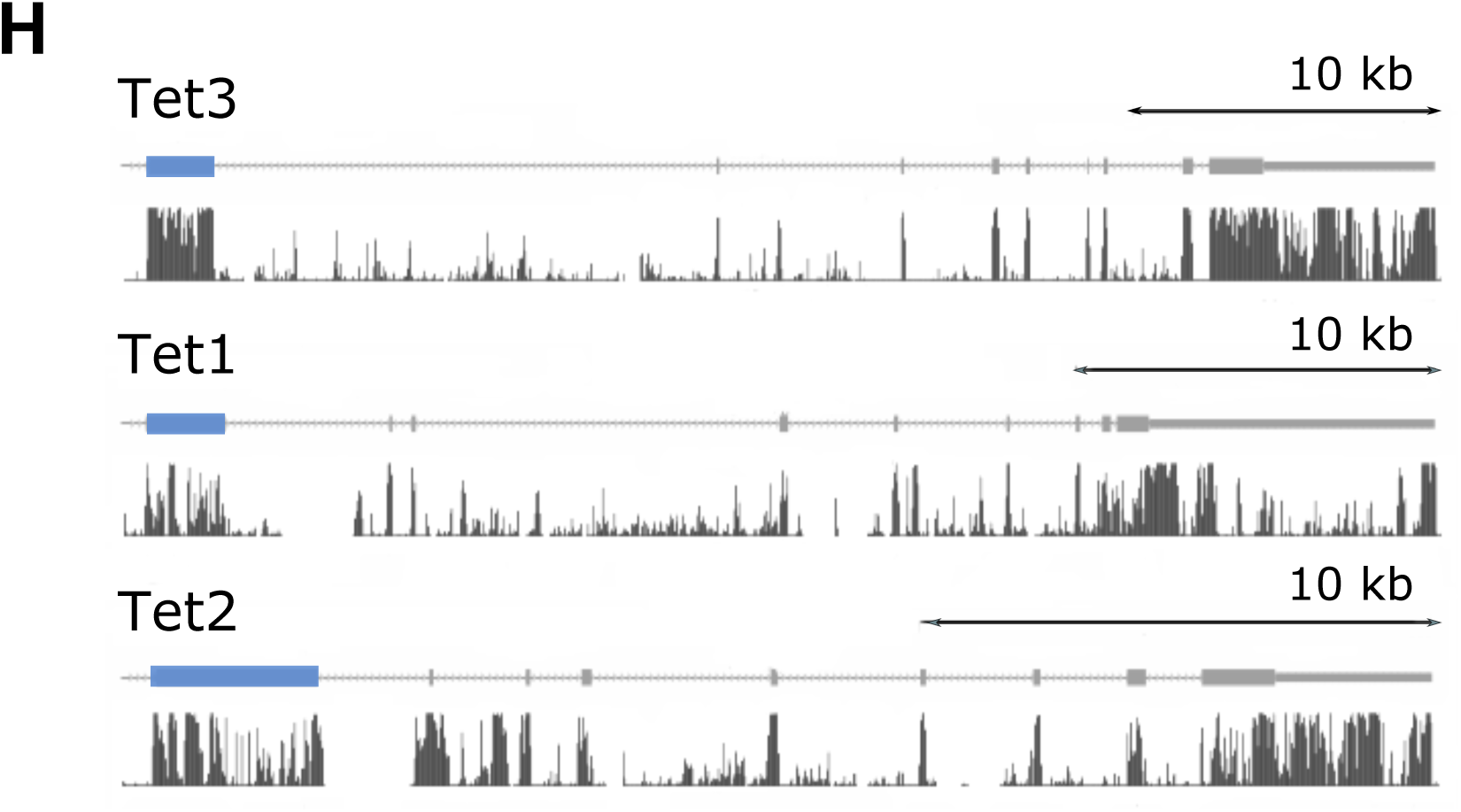
A) Expression changes of a group of genes differentially regulated by TET3 for cells that either overexpress the first half of the protein (TET3_trunc_) or the second half (CD, catalytic domain), analysed by qPCR and shown as fold changes of TET3 variant expressing versus control cells. TET3_trunc_ increases expression whereas the CD does not. B) Percentage of CD40 (trophoblast marker) positive cells at the end point of a transdifferentiation experiment. Trophoblast stem (TS) cells were used as a positive control for CD40 staining. Oct4 KO cells were used as a model for facilitated trans-differentiation (Niwa et al. 2000, 2005). Control cells are GFP negative cells from the Tet3_trunc_ transgenic line. TET3_trunc_ expressing cells show a significantly higher proportion of transdifferentiated cells compared to control cells as determined by Chi-squared test (**** p<0.0001). C) Heatmap of TET3 ChIP-seq peaks identified using MACS peak calling. Peaks identified for each individual sample were combined and reads in peaks quantified in all samples. Every row represents one peak. Individual replicates are shown and grouped as denoted by horizontal lines below. Signal in peaks was extremely similar between TET3, TET3_CXXC_ and TET3_trunc_. D) Quantitative comparison of TET3 interaction partners by Stable Isotope Labelling with Amino Acids in Cell culture (SILAC). For every protein the number of peptides found in the TET3 sample was divided by the number of peptides found in the TET3_trunc_ samples, and normalised for the ratio of found TET3 peptides. The list was manually curated to exclude actins, keratins and ribosomal proteins. The high log_2_ ratios for JIP3 and OGT indicate that their interactions with TET3 are lost upon deletion of the catalytic domain (see also Table S4). E) Cells were transiently transfected with siRNA against OGT and then TET3 expression was induced. Cells were sorted for TET3 and a group of genes differentially regulated by TET3 analysed by qPCR. Shown are expression fold changes of TET3 positive versus TET3 negative cells for cells treated with siRNA against OGT or untransfected cells (NT). Genes normally upregulated upon TET3 expression are still upregulated in the absence of OGT. F) Expression changes of a group of genes differentially regulated by TET3 for cells that either overexpress TET3 (oocyte isoform) or TET3_CXXC_, analysed by qPCR and shown as log_2_ fold changes of TET3 variant expressing versus control cells. TET3 and TET3_CXXC_ have a similar effect on expression. G) Expression changes of a group of genes differentially regulated by TET3 for cells that either overexpress TET3, TET3_trunc_, exon 4 only or GFP (see Fig. 5B for constructs), analysed by qPCR and shown as fold change between TET3 variant expressing versus control cells. Exon 4 by itself induces the similar expression changes as TET3 and TET3_trunc_. H) Zoomed out view of Figure 5H: Evolutionary conservation across placental mammals of Tet3 (exon 4 to end) and the corresponding regions of other Tet family members was measured using PhastCons (Siepel et al. 2005). Exons are represented as grey boxes with exon 4 and corresponding exons highlighted in blue. Conservation scores between 0 and 1 are represented by the height of the black bars.

## References

Adachi K, Nikaido I, Ohta H, Ohtsuka S, Ura H, Kadota M, Wakayama T, Ueda HR, Niwa H. 2013. Context-Dependent Wiring of Sox2 Regulatory Networks for Self-Renewal of Embryonic and Trophoblast Stem Cells. Mol Cell 52: 380–392.

Allis CD, Jenuwein T. 2016. The molecular hallmarks of epigenetic control. Nat Rev Genet 17: 487–500.

Amouroux R, Nashun B, Shirane K, Nakagawa S, Hill PW, D’Souza Z, Nakayama M, Matsuda M, Turp A, Ndjetehe E, et al. 2016. De novo DNA methylation drives 5hmC accumulation in mouse zygotes. Nat Cell Biol 18: 225–233.

Apostolou E, Hochedlinger K. 2013. Chromatin dynamics during cellular reprogramming. Nature 502: 462–471.

Barrero MJ, Boué S, Izpisúa Belmonte JC. 2010. Epigenetic Mechanisms that Regulate Cell Identity. Cell Stem Cell 7: 565–570.

Borsos M, Torres-Padilla M-E. 2016. Building up the nucleus: nuclear organization in the establishment of totipotency and pluripotency during mammalian development. Genes Dev 30: 611–621.

Branco MR, Ficz G, Reik W. 2012. Uncovering the role of 5-hydroxymethylcytosine in the epigenome. Nat Rev Genet 13: 7–13.

Buenrostro JD, Giresi PG, Zaba LC, Chang HY, Greenleaf WJ. 2013. Transposition of native chromatin for fast and sensitive epigenomic profiling of open chromatin, DNA-binding proteins and nucleosome position. Nat Methods 10: 1213–1218.

Cambuli F, Murray A, Dean W, Dudzinska D, Krueger F, Andrews S, Senner CE, Cook SJ, Hemberger M. 2014. Epigenetic memory of the first cell fate decision prevents complete ES cell reprogramming into trophoblast. Nat Commun 5.

Chen Q, Chen Y, Bian C, Fujiki R, Yu X. 2013. TET2 promotes histone O-GlcNAcylation during gene transcription. Nature 493: 561–564.

Chen T, Dent SYR. 2014. Chromatin modifiers and remodellers: regulators of cellular differentiation. Nat Rev Genet 15: 93–106.

Clift D, Schuh M. 2013. Restarting life: fertilization and the transition from meiosis to mitosis. Nat Rev Mol Cell Biol 14: 549–562.

Creyghton MP, Cheng AW, Welstead GG, Kooistra T, Carey BW, Steine EJ, Hanna J, Lodato MA, Frampton GM, Sharp PA, et al. 2010. Histone H3K27ac separates active from poised enhancers and predicts developmental state. Proc Natl Acad Sci 107: 21931–21936.

Dai H-Q, Wang B-A, Yang L, Chen J-J, Zhu G-C, Sun M-L, Ge H, Wang R, Chapman DL, Tang F, et al. 2016. TET-mediated DNA demethylation controls gastrulation by regulating Lefty–Nodal signalling. Nature 538: 528–532.

Deplus R, Delatte B, Schwinn MK, Defrance M, Méndez J, Murphy N, Dawson MA, Volkmar M, Putmans P, Calonne E, et al. 2013. TET2 and TET3 regulate GlcNAcylation and H3K4 methylation through OGT and SET1/COMPASS. EMBO J 32: 645–655.

Eckersley-Maslin MA, Svensson V, Krueger C, Stubbs TM, Giehr P, Krueger F, Miragaia RJ, Kyriakopoulos C, Berrens RV, Milagre I, et al. 2016. MERVL/Zscan4 Network Activation Results in Transient Genome-wide DNA Demethylation of mESCs. Cell Rep 17: 179–192.

Evsikov AV, Vries WN de, Peaston AE, Radford EE, Fancher KS, Chen FH, Blake JA, Bult CJ, Latham KE, Solter D, et al. 2004. Systems biology of the 2-cell mouse embryo. Cytogenet Genome Res 105: 240–250.

Gu T-P, Guo F, Yang H, Wu H-P, Xu G-F, Liu W, Xie Z-G, Shi L, He X, Jin S, et al. 2011. The role of Tet3 DNA dioxygenase in epigenetic reprogramming by oocytes. Nature 477: 606–610.

Guo F, Li X, Liang D, Li T, Zhu P, Guo H, Wu X, Wen L, Gu T-P, Hu B, et al. 2014. Active and Passive Demethylation of Male and Female Pronuclear DNA in the Mammalian Zygote. Cell Stem Cell 15: 447–458.

Guzman-Ayala M, Sachs M, Koh FM, Onodera C, Bulut-Karslioglu A, Lin C-J, Wong P, Nitta R, Song JS, Ramalho-Santos M. 2015. Chd1 is essential for the high transcriptional output and rapid growth of the mouse epiblast. Development 142: 118–127.

Hahn MA, Qiu R, Wu X, Li AX, Zhang H, Wang J, Jui J, Jin S-G, Jiang Y, Pfeifer GP, et al. 2013. Dynamics of 5-Hydroxymethylcytosine and Chromatin Marks in Mammalian Neurogenesis. Cell Rep 3: 291–300.

He Y-F, Li B-Z, Li Z, Liu P, Wang Y, Tang Q, Ding J, Jia Y, Chen Z, Li L, et al. 2011. Tet-Mediated Formation of 5-Carboxylcytosine and Its Excision by TDG in Mammalian DNA. Science 333: 1303–1307.

Hosseini SM, Dufort I, Nieminen J, Moulavi F, Ghanaei HR, Hajian M, Jafarpour F, Forouzanfar M, Gourbai H, Shahverdi AH, et al. 2016. Epigenetic modification with trichostatin A does not correct specific errors of somatic cell nuclear transfer at the transcriptomic level; highlighting the non-random nature of oocyte-mediated reprogramming errors. BMC Genomics 17: 16.

Iqbal K, Jin S-G, Pfeifer GP, Szabó PE. 2011. Reprogramming of the paternal genome upon fertilization involves genome-wide oxidation of 5-methylcytosine. Proc Natl Acad Sci U S A 108: 3642–3647.

Ito R, Katsura S, Shimada H, Tsuchiya H, Hada M, Okumura T, Sugawara A, Yokoyama A. 2014. TET3–OGT interaction increases the stability and the presence of OGT in chromatin. Genes Cells 19: 52–65.

Ito S, D’Alessio AC, Taranova OV, Hong K, Sowers LC, Zhang Y. 2010. Role of Tet proteins in 5mC to 5hmC conversion, ES-cell self-renewal and inner cell mass specification. Nature 466: 1129–1133.

Ito S, Shen L, Dai Q, Wu SC, Collins LB, Swenberg JA, He C, Zhang Y. 2011. Tet Proteins Can Convert 5-Methylcytosine to 5-Formylcytosine and 5-Carboxylcytosine. Science 333: 1300–1303.

Jao CY, Salic A. 2008. Exploring RNA transcription and turnover in vivo by using click chemistry. Proc Natl Acad Sci U S A 105: 15779–15784.

Jin S-G, Zhang Z-M, Dunwell TL, Harter MR, Wu X, Johnson J, Li Z, Liu J, Szabó PE, Lu Q, et al. 2016. Tet3 reads 5-carboxylcytosine through its CXXC domain and is a potential guardian against neurodegeneration. Cell Rep 14: 493–505.

Kang J, Lienhard M, Pastor WA, Chawla A, Novotny M, Tsagaratou A, Lasken RS, Thompson EC, Surani MA,Koralov SB, et al. 2015. Simultaneous deletion of the methylcytosine oxidases Tet1 and Tet3 increases transcriptome variability in early embryogenesis. Proc Natl Acad Sci 112: E4236–E4245.

Kigami D, Minami N, Takayama H, Imai H. 2003. MuERV-L is one of the earliest transcribed genes in mouse one-cell embryos. Biol Reprod 68: 651–654.

Kim A, Pyykko I. 2011. Size matters: versatile use of PiggyBac. Mol Cell Biochem 354: 301–309.

Kohli RM, Zhang Y. 2013. TET enzymes, TDG and the dynamics of DNA demethylation. Nature 502: 472–479.

Krishnakumar R, Blelloch RH. 2013. Epigenetics of cellular reprogramming. Curr Opin Genet Dev 23: 548–555.

Lee HJ, Hore TA, Reik W. 2014. Reprogramming the methylome: erasing memory and creating diversity. Cell Stem Cell 14: 710–719.

Li L, Zheng P, Dean J. 2010. Maternal control of early mouse development. Dev Camb Engl 137: 859–870.

Lian H, Li W-B, Jin W-L, Lian H, Li W-B, Jin W-L. 2016. The emerging insights into catalytic or non-catalytic roles of TET proteins in tumors and neural development. Oncotarget 7: 64512–64525.

Long HK, Blackledge NP, Klose RJ. 2013. ZF-CxxC domain-containing proteins, CpG islands and the chromatin connection. Biochem Soc Trans 41: 727–740.

Love MI, Huber W, Anders S. 2014. Moderated estimation of fold change and dispersion for RNA-seq data with DESeq2. Genome Biol 15: 550.

Macfarlan TS, Gifford WD, Agarwal S, Driscoll S, Lettieri K, Wang J, Andrews SE, Franco L, Rosenfeld MG, Ren B, et al. 2011. Endogenous retroviruses and neighboring genes are coordinately repressed by LSD1/KDM1A. Genes Dev 25: 594–607.

Milagre I, Stubbs TM, King MR, Spindel J, Santos F, Krueger F, Bachman M, Segonds-Pichon A, Balasubramanian S, Andrews SR, et al. 2017. Gender Differences in Global but Not Targeted Demethylation in iPSC Reprogramming. Cell Rep 18: 1079–1089.

Mohammed H, D’Santos C, Serandour AA, Ali HR, Brown GD, Atkins A, Rueda OM, Holmes KA, Theodorou V, Robinson JLL, et al. 2013. Endogenous Purification Reveals GREB1 as a Key Estrogen Receptor Regulatory Factor. Cell Rep 3: 342–349.

Mohammed H, Taylor C, Brown GD, Papachristou EK, Carroll JS, D’Santos CS. 2016. Rapid immunoprecipitation mass spectrometry of endogenous proteins (RIME) for analysis of chromatin complexes. Nat Protoc 11: 316–326.

Ng RK, Dean W, Dawson C, Lucifero D, Madeja Z, Reik W, Hemberger M. 2008. Epigenetic restriction of embryonic cell lineage fate by methylation of Elf5. Nat Cell Biol 10: 1280–1290.

Ni K, Dansranjavin T, Rogenhofer N, Oeztuerk N, Deuker J, Bergmann M, Schuppe H-C, Wagenlehner F, Weidner W, Steger K, et al. 2016. TET enzymes are successively expressed during human spermatogenesis and their expression level is pivotal for male fertility. Hum Reprod 31: 1411– 1424.

Niwa H, Miyazaki J, Smith AG. 2000. Quantitative expression of Oct-3/4 defines differentiation, dedifferentiation or self-renewal of ES cells. Nat Genet 24: 372–376.

Niwa H, Toyooka Y, Shimosato D, Strumpf D, Takahashi K, Yagi R, Rossant J. 2005. Interaction between Oct3/4 and Cdx2 Determines Trophectoderm Differentiation. Cell 123: 917–929.

Pastor WA, Aravind L, Rao A. 2013. TETonic shift: biological roles of TET proteins in DNA demethylation and transcription. Nat Rev Mol Cell Biol 14: 341–356.

Peaston AE, Evsikov AV, Graber JH, de Vries WN, Holbrook AE, Solter D, Knowles BB. 2004. Retrotransposons Regulate Host Genes in Mouse Oocytes and Preimplantation Embryos. Dev Cell 7: 597–606.

Peat JR, Dean W, Clark SJ, Krueger F, Smallwood SA, Ficz G, Kim JK, Marioni JC, Hore TA, Reik W. 2014. Genome-wide Bisulfite Sequencing in Zygotes Identifies Demethylation Targets and Maps the Contribution of TET3 Oxidation. Cell Rep 9: 1990–2000.

Percharde M, Bulut-Karslioglu A, Ramalho-Santos M. 2017a. Hypertranscription in Development, Stem Cells, and Regeneration. Dev Cell 40: 9–21.

Percharde M, Wong P, Ramalho-Santos M. 2017b. Global Hypertranscription in the Mouse Embryonic Germline. Cell Rep 19: 1987–1996.

Perera A, Eisen D, Wagner M, Laube SK, Künzel AF, Koch S, Steinbacher J, Schulze E, Splith V, Mittermeier N, et al. 2015. TET3 Is Recruited by REST for Context-Specific Hydroxymethylation and Induction of Gene Expression. Cell Rep 11: 283–294.

Polo JM, Anderssen E, Walsh RM, Schwarz BA, Nefzger CM, Lim SM, Borkent M, Apostolou E, Alaei S, Cloutier J, et al. 2012. A Molecular Roadmap of Reprogramming Somatic Cells into iPS Cells. Cell 151: 1617–1632.

Rahl PB, Lin CY, Seila AC, Flynn RA, McCuine S, Burge CB, Sharp PA, Young RA. 2010. c-Myc Regulates Transcriptional Pause Release. Cell 141: 432–445.

Rasmussen KD, Helin K. 2016. Role of TET enzymes in DNA methylation, development, and cancer. Genes Dev 30: 733–750.

Rice P, Longden I, Bleasby A. 2000. EMBOSS: The European Molecular Biology Open Software Suite. Trends Genet 16: 276–277.

Rossant J. 2008. Stem Cells and Early Lineage Development. Cell 132: 527–531.

Rugg-Gunn PJ, Cox BJ, Lanner F, Sharma P, Ignatchenko V, McDonald ACH, Garner J, Gramolini AO, Rossant J, Kislinger T. 2012. Cell-Surface Proteomics Identifies Lineage-Specific Markers of Embryo-Derived Stem Cells. Dev Cell 22: 887–901.

Santos F, Peat J, Burgess H, Rada C, Reik W, Dean W. 2013. Active demethylation in mouse zygotes involves cytosine deamination and base excision repair. Epigenetics Chromatin 6: 39.

Schmidt D, Wilson MD, Spyrou C, Brown GD, Hadfield J, Odom DT. 2009. ChIP-seq: Using high-throughput sequencing to discover protein–DNA interactions. Methods 48: 240–248.

Seisenberger S, Andrews S, Krueger F, Arand J, Walter J, Santos F, Popp C, Thienpont B, Dean W, Reik W. 2012. The Dynamics of Genome-wide DNA Methylation Reprogramming in Mouse Primordial Germ Cells. Mol Cell 48: 849–862.

Shen L, Inoue A, He J, Liu Y, Lu F, Zhang Y. 2014. Tet3 and DNA Replication Mediate Demethylation of Both the Maternal and Paternal Genomes in Mouse Zygotes. Cell Stem Cell 15: 459–470.

Siepel A, Bejerano G, Pedersen JS, Hinrichs AS, Hou M, Rosenbloom K, Clawson H, Spieth J, Hillier LW,Richards S, et al. 2005. Evolutionarily conserved elements in vertebrate, insect, worm, and yeast genomes. Genome Res 15: 1034–1050.

Skene PJ, Hernandez AE, Groudine M, Henikoff S. 2014. The nucleosomal barrier to promoter escape by RNA polymerase II is overcome by the chromatin remodeler Chd1. eLife 3: e02042.

Smallwood SA, Tomizawa S-I, Krueger F, Ruf N, Carli N, Segonds-Pichon A, Sato S, Hata K, Andrews SR, Kelsey G. 2011. Dynamic CpG island methylation landscape in oocytes and preimplantation embryos. Nat Genet 43: 811–814.

Soufi A, Dalton S. 2016. Cycling through developmental decisions: how cell cycle dynamics control pluripotency, differentiation and reprogramming. Development 143: 4301–4311.

Tahiliani M, Koh KP, Shen Y, Pastor WA, Bandukwala H, Brudno Y, Agarwal S, Iyer LM, Liu DR, Aravind L, et al. 2009. Conversion of 5-Methylcytosine to 5-Hydroxymethylcytosine in Mammalian DNA by MLL Partner TET1. Science 324: 930–935.

Tan L, Shi YG. 2012. Tet family proteins and 5-hydroxymethylcytosine in development and disease. Development 139: 1895–1902.

Tsukada Y, Akiyama T, Nakayama KI. 2015. Maternal TET3 is dispensable for embryonic development but is required for neonatal growth. Sci Rep 5: 15876.

Veillard A-C, Marks H, Bernardo AS, Jouneau L, Laloë D, Boulanger L, Kaan A, Brochard V, Tosolini M, Pedersen R, et al. 2014. Stable Methylation at Promoters Distinguishes Epiblast Stem Cells from Embryonic Stem Cells and the In Vivo Epiblasts. Stem Cells Dev 23: 2014–2029.

Vella P, Scelfo A, Jammula S, Chiacchiera F, Williams K, Cuomo A, Roberto A, Christensen J, Bonaldi T, Helin K, et al. 2013. Tet Proteins Connect the O-Linked N-acetylglucosamine Transferase Ogt to Chromatin in Embryonic Stem Cells. Mol Cell 49: 645–656.

Veselovska L, Smallwood SA, Saadeh H, Stewart KR, Krueger F, Maupetit-Méhouas S, Arnaud P, Tomizawa S, Andrews S, Kelsey G. 2015. Deep sequencing and de novo assembly of the mouse oocyte transcriptome define the contribution of transcription to the DNA methylation landscape. Genome Biol 16: 209.

Wossidlo M, Nakamura T, Lepikhov K, Marques CJ, Zakhartchenko V, Boiani M, Arand J, Nakano T, Reik W, Walter J. 2011. 5-Hydroxymethylcytosine in the mammalian zygote is linked with epigenetic reprogramming. Nat Commun 2: 241.

Xu Y, Wu F, Tan L, Kong L, Xiong L, Deng J, Barbera A, Zheng L, Zhang H, Huang S, et al. 2011. Genome-wide Regulation of 5hmC, 5mC and Gene Expression by Tet1 Hydroxylase in Mouse Embryonic Stem Cells. Mol Cell 42: 451–464.

Xu Y, Xu C, Kato A, Tempel W, Abreu JG, Bian C, Hu Y, Hu D, Zhao B, Cerovina T, et al. 2012. Tet3 CXXC Domain and Dioxygenase Activity Cooperatively Regulate Key Genes for Xenopus Eye and Neural Development. Cell 151: 1200–1213.

Xue Z, Huang K, Cai C, Cai L, Jiang C, Feng Y, Liu Z, Zeng Q, Cheng L, Sun YE, et al. 2013. Genetic programs in human and mouse early embryos revealed by single-cell RNA?sequencing. Nature 500: 593–597.

Yu C, Zhang Y-L, Pan W-W, Li X-M, Wang Z-W, Ge Z-J, Zhou J-J, Cang Y, Tong C, Sun Q-Y, et al. 2013. CRL4 Complex Regulates Mammalian Oocyte Survival and Reprogramming by Activation of TET Proteins. Science 342: 1518–1521.

Yue F, Cheng Y, Breschi A, Vierstra J, Wu W, Ryba T, Sandstrom R, Ma Z, Davis C, Pope BD, et al. 2014. A comparative encyclopedia of DNA elements in the mouse genome. Nature 515: 355–364.

Zhang Q, Liu X, Gao W, Li P, Hou J, Li J, Wong J. 2014. Differential Regulation of Ten-Eleven Translocation Family of Dioxygenases by O-Linked β-N-Acetylglucosamine Transferase OGT. J Biol Chem jbc.M113.524140.

Zhang Y, Liu T, Meyer CA, Eeckhoute J, Johnson DS, Bernstein BE, Nusbaum C, Myers RM, Brown M, Li W, et al. 2008. Model-based Analysis of ChIP-Seq (MACS). Genome Biol 9: R137.

